# A Systems Biology Approach to Unveil Shared Therapeutic Targets and Pathological Pathways Across Major Human Cancers

**DOI:** 10.1101/2025.05.14.653945

**Authors:** Aftab Alam, Mohd Faizan Siddiqui, Rifat Hamoudi, Uday Kishore, Maria Fernandez Cabezudo, Basel K. Al-Ramadi

## Abstract

Cancer is a leading global cause of mortality, responsible for nearly 10 million deaths annually, with breast, lung, colorectal, and prostate cancers among the most prevalent. Despite extensive research on individual cancer types, identifying shared molecular signatures could unlock pan-cancer diagnostic tools and therapeutic targets. This study leveraged RNA-seq data from The Cancer Genome Atlas (TCGA) to analyze our selected cancer types (SCTs)—breast, lung, colorectal, and prostate, employing a multi-step bioinformatics pipeline. Differentially expressed genes (DEGs) between tumors and normal tissues were identified and validated using an Elastic Net regression model. Weighted Gene Co-expression Network Analysis (WGCNA) revealed highly correlated gene modules linked to clinical traits, pinpointing 179 shared signature genes across the SCTs. Protein-protein interaction (PPI) network and clustering analyses further refined these to 26 hub genes, enriched in cancer hallmark pathways. Nine hub genes—*KIF18B, RRM2, MYBL2, IQGAP3, TPX2, SLC7A11, RHPN1, HJURP*, and *SKA3*—stood out due to consistent upregulation in metastatic tumors (breast, colorectal, prostate) and their high expression across more than 18 different cancer types, suggesting roles as oncogenes, prognostic markers, or therapeutic targets. The expression patterns of hub genes were further validated across larger cancer patient cohorts of SCTs, confirming relevance across multiple datasets, and their prognostic significance was assessed by their influence on overall survival (OS). Notably, these hub genes also correlated with immune-related functions, potentially influencing tumor microenvironment modulation. This integrative approach provides a strong framework for identifying cross-cancer potential biomarkers, advancing pan-cancer insights, and supporting improved diagnosis, prognosis, and therapy development.

## 1. Introduction

Cancer remains a major global public health challenge, driven by aging populations, lifestyle factors, infections, and environmental exposures. It disproportionately affects low- and middle-income countries (LMICs), which face limited access to prevention, diagnosis, and treatment. In 2022, an estimated 20 million new cases and 9.7 million cancer deaths occurred globally, with nearly 70% of deaths in LMICs. By 2050, cancer cases are projected to rise by 87.5%, placing even more pressure on health systems, individuals, and communities [1]. According to the <u>World Health Organization</u> (WHO), the most common cancers are breast, lung, colon, and prostate cancers, which were selected for this study as they originate from diverse tissue types, providing a robust framework and rationale.

Breast cancer is the most common cancer among women, making up 12.5% of all new cancer cases globally. In 2022, 2.3 million women were diagnosed with breast cancer, resulting in 670,000 deaths [2]. Lung cancer affects ∼2.5 million people each year and claims 1.8 million lives annually [3]. Colorectal cancer, with >1.9 million diagnoses in 2022, is the third most common cancer globally, responsible for more than 900,000 deaths each year. Prostate cancer affects 1 in 8 men during their lifetime and resulted in 1.4 million diagnoses and 375,000 deaths in 2020 [4].

To reduce the cancer burden, early detection through screening and quick treatment improves survival. Biomarker detection plays a critical role in this process by identifying cancer at an early stage, confirming its type, and guiding treatment choices. Specific biomarkers can also help predict how aggressive the cancer is and how it will respond to treatments, enabling personalized therapy. High-throughput gene expression technologies offer extensive genetic insights into cancer types, revealing crucial changes in disease progression [5–8]. By leveraging genomics, epigenomics, and transcriptomics data available from online databases, it is possible to uncover potentially novel biomarkers associated with cancer [9–12]. Recently, machine learning models such as support vector machines (SVMs), random forests, linear models, and Elastic Net models have emerged as effective tools for predictive gene signatures [13–15]. These approaches hold promise for advancing our understanding and diagnosis of cancer.

However, despite significant advances in high-throughput technologies and data integration approaches, identifying common biomarkers remains challenging due to cancer heterogeneity. Most existing studies focus on individual cancer types, limiting our understanding of common oncogenic processes that could be targeted across cancers. This creates a critical research gap in identifying pan-cancer biomarkers and immune modulators that could lead to more universally applicable therapeutic strategies.

This study addresses the gap in understanding cancer-specific molecular signatures by leveraging TCGA datasets to identify common differentially expressed genes (DEGs) across breast, lung, colon, and prostate cancers—referred to as selected cancer types (SCTs), which are among the most prevalent cancers. We then applied Weighted Gene Co-expression Network Analysis (WGCNA) to identify modules of co-expressed genes [16,17]. Subsequently, a protein-protein interaction (PPI) network was then constructed to evaluate their topological significance within the global interactome. This network analysis provided insights into the nature and strength of gene interactions, while functional clustering helped uncover signature genes (hub genes) and their roles in dysregulated cancer-related pathways. Additionally, we explored the role of the immune system in cancer by examining the relationship between the identified hub genes and immune-related features, aiming to better understand their involvement in modulating the immune landscape.

This cross-cancer transcriptomic analysis reveals conserved expression patterns of DEGs, identifies common potential biomarkers, and highlights key biological pathways across selected cancer types (SCTs). By combining WGCNA and PPI network analysis, we enhanced the detection of biologically significant hub genes within co-expression modules and their protein-level interactions. Moreover, by linking these hub genes to immune-related features, we offer novel insights into their potential roles in modulating the tumor immune microenvironment across SCTs. These insights contribute to improved diagnostic accuracy, uncover opportunities for drug repurposing, and support the development of broad-spectrum or personalized therapies. This work provides in silico insights that can guide experimental research, leading to more effective targeted therapies that could reduce cancer-related morbidity and mortality. However, experimental validation of these findings is needed to confirm their relevance and further explore their potential clinical applications.

## 2. Materials and Methods

A schematic diagram describing the workflow utilized for the analysis is shown in **Fig. 1**.

**Figure 1:**
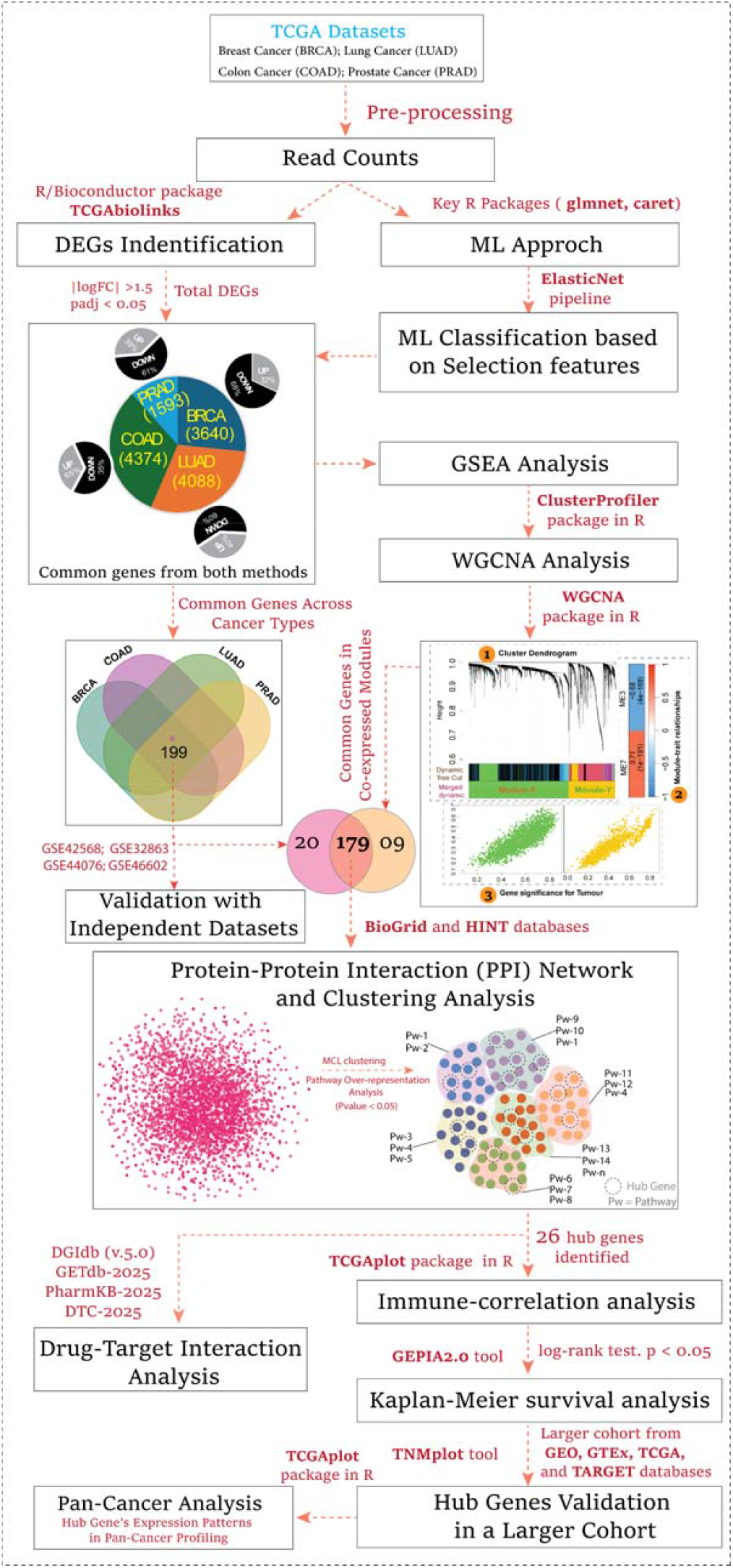
Overview of the study workflow, including key steps such as data collection, preprocessing, analysis, and validation

### 2.1. Data Collection and Differential Expression Analysis

Our study analyzed four selected cancer types (SCTs) from The Cancer Genome Atlas (TCGA), a program developed by the National Institutes of Health (NIH), USA. The dataset included: breast invasive carcinoma (BRCA), lung adenocarcinoma (LUAD), colon adenocarcinoma (COAD), and prostate adenocarcinoma (PRAD). RNA expression data for cancers were obtained from the TCGA biolinks package in R. Differential expression analysis between ‘Primary Tumor’ and ‘Solid Tissue Normal’ samples in cancers was conducted using the limma package, which is commonly used for RNA-seq data analysis. Raw gene count data were normalized using the TMM (Trimmed Mean of M-values) method to account for differences in library sizes and sequencing depth across samples. The normalization process was implemented using the calcNormFactors() function, followed by the **voom** transformation, which transformed the count data into log-counts per million (log-CPM) and estimated the mean-variance relationship for each gene.

Next, a design matrix was built using the model.matrix() function based on the condition variable “definition” (e.g., Tumor vs. Normal). The analysis included 1,111 tumor and 113 normal samples for BRCA, 539 tumor and 59 normal samples for LUAD, 481 tumor and 41 normal samples for COAD, and 501 tumor and 52 normal samples for PRAD. Differential expression analysis was performed using lmFit() and eBayes() from the **limma** package. DEGs were identified based on an absolute fold change > 1.5 and an adjusted p-value < 0.05, using the Benjamini-Hochberg (BH) method for multiple testing correction. We also identified a set of commonly dysregulated genes shared across selected cancer types (SCTs), exhibiting consistent expression patterns [**Suppl. Fig. 1–2**]. This suggests their potential involvement in core biological processes or conserved oncogenic pathways.

### 2.2. Machine Learning Classification

We explored machine learning to classify unseen samples as tumor or non-tumor, starting with a simple linear model with feature selection, followed by an Elastic Net model [18]. Here, we used expression data that has already been normalized, along with clinical features such as tumor vs. normal across each SCT individually. For each SCT, we split 75% of the samples into a training set and 25% into a test set using stratified sampling with the createDataPartition() function from the caret package to maintain class balance. We then trained a regularized logistic regression model (Elastic Net with α = 0.5) using cross-validation via the cv.glmnet() function from the glmnet package. Model performance was assessed by a confusion matrix, comparing predictions to actual values (true positives, true negatives, false positives, false negatives). Then we analyzed the genes selected by Elastic Net, focusing on those with non-zero coefficients as key predictors. Next, we clustered all samples (Tumor and Normal) using hierarchical clustering. To assess the reliability of our DEG identification, we performed cross-validation with Elastic Net and selected genes predicted by both the Limma and Elastic Net models [**Suppl. Fig. 3**].

### 2.3. Gene Set Enrichment Analysis (GSEA)

All DEGs from SCTs were further analyzed using GSEA with Reactome pathways through the gsePathway function from the clusterProfiler package in R. We set the threshold for pathway selection with pvalueCutoff < 0.05, minGSSize = 2, and maxGSSize = 500. All the pathways were mapped over the Reactome hierarchy map [19] obtained from the Reactome database (<u>Pathways hierarchy relationship data</u>). In this network map, each node represents a biological pathway, and edges illustrate hierarchical or functional relationships between them. Node size corresponds to the number of genes in the pathway (gene set size), while node color reflects the normalized enrichment score (NES), with red indicating positively enriched pathways and green indicating negatively enriched ones, representing the direction and magnitude of enrichment. The border color of each node denotes the statistical significance (p-value), using a gradient from dark to light to show decreasing significance—darker borders indicate higher statistical confidence. This integrated visualization highlights both the strength and reliability of pathway enrichment, as well as their biological interconnections.

### 2.4. Validation with Independent Datasets

We further validated the commonly-expressed genes across SCTs using independent datasets to ensure robustness, reliability, and generalizability of our findings. We used GEO datasets, including BRCA (GSE42568, 104 tumors, 17 normals) [20], LUAD (GSE32863, 58 tumors, 58 normals) [21], COAD (GSE44076, 98 tumors, 50 normals) [22], and PRAD (GSE46602, 36 tumors, 14 normals) [23]. In this validation, we focused on the directionality of gene expression changes—whether upregulated or downregulated (rather than significance level), to capture consistent expression patterns across datasets. Further, we compute the Pearson correlation (r) and p-value between TCGA and GEO gene expression values of all 199 common DEGs (for both tumor and normal samples) to assess the reproducibility quantitatively, validate the biological relevance, and demonstrate the robustness of our observed gene expression patterns across independent datasets.

### 2.5. Weighted Gene Correlation Network Analysis (WGCNA)

Traditional gene expression analysis identifies DEGs between tumor and normal datasets, treating each gene independently and overlooking the complex interconnections within the transcriptome. This limits its ability to identify key drivers of disease affected by diverse factors influencing gene expression profiles. In contrast to random networks, scale-free networks exhibit a notable abundance of highly connected nodes known as “hubs” [16,17]. These hubs are central within their networks, reflecting specific functional roles. The WGCNA uses scale-free networks to identify gene relationships, detecting modules of highly correlated genes and key hub genes that contribute to phenotypic traits.

In this study, a scale-free network using gene expression profiles of DEGs was constructed using the WGCNA_V1.72.5 package in R [24] to build a weighted correlation network from tumor and normal SCT datasets. Gene expression matrices were transformed using the Pearson correlation coefficient to create similarity matrices for paired genes, which were then converted into adjacency matrices using a soft-threshold (power values) to highlight significant gene connections and filter out weak correlations. The adjacency matrix was converted to a topological overlap measure (TOM) to assess gene interaction strengths. Genes were hierarchically clustered based on TOM values, and modules were identified using the DynamicTreeCut function; modules with high similarity scores were merged. Additionally, module eigengenes (MEs) represented each module’s expression profile, and module membership (MM) was defined as the correlation between MEs and gene expression. Further, we checked the relationship between module genes and specific traits by analyzing the distribution of their correlation values.

### 2.6. PPI Network, Clustering, and Hub Gene Identification

We constructed a PPI network of the 179 identified common genes to examine their topology within the global interactome. Analyzing the topological parameters of a PPI network can provide valuable insights into the quality and nature of interactions, shedding light on the connections and relationships between nodes (i.e., genes) in the network. To build the PPI network, we integrated entries from BioGrid (v.4.4.236)) [25] and HINT (High-quality Interactomes, version 2024-06) [26] to create a comprehensive dataset of documented human-human (Homo sapiens) PPIs. Although HINT already compiles high-quality PPIs from eight interactome resources, we combined these datasets to create a more robust network and maximize interaction coverage for our genes of interest. Next, clustering was performed using the unsupervised MCL (Markov Cluster Algorithm) method [34] to identify functional clusters based on stochastic flow in the PPI network by setting parameters with a degree cut-off of ≥ 3, 1000 iterations, and an inflation parameter of ≥ 2. The identified modules revealed key biological processes and pathways.

### 2.7. Over-representation analysis of Functional Modules

After identifying clusters in the PPI network, we performed functional pathway analysis for each cluster with hub genes and their partners using the Enrichr web tool [27] that can be found at https://maayanlab.cloud/Enrichr/. Enrichr allows users to submit lists of genes (human or mouse) for comparison against various gene set libraries, such as MSigDB [28], KEGG [29], <u>Elsevier pathway collection (EPC)</u> and <u>Reactome database</u> [30]. We integrate all databases for pathway analysis to ensure comprehensive coverage, as some pathways may be present in one database but not others for the query gene set, providing more robust insights into the clusters. These databases are reliable due to their rigorous curation, comprehensive pathway coverage, and regular updates. We set the adjusted p-value threshold to < 0.05 and the minimum gene set size to ≥ 2.

### 2.8 Correlation analysis of the Key Genes with Immune-related Genes

We performed correlation analysis between the 26 hub gene sets (from SCTs) and immune-related genes, immune cell ratios, and immune score using TCGAplot [31] package in R, designed for visualizing and analyzing TCGA data. The goal was to identify significant patterns and associations to better understand the immune landscape in SCTs and the role of the key gene set in modulating immune responses.

### 2.9 Survival Analysis of the Key Genes in Functional Modules

To evaluate the prognostic values of the 26 hub genes, we carried out Kaplan-Meier survival analysis for overall survival (OS) using GEPIA2.0 [32]. We selected the multiple dataset option for the SCTs. For each key gene, the group cutoff was set at the quartile, and samples were divided into high-expression and low-expression cohorts based on whether their expression levels were above or below the threshold. Statistical significance was assessed using the log-rank test, with p<0.05 considered statistically significant.

The total number of patients for KM analysis was: BRCA (535 high, 534 low), COAD (134 high, 135 low), LUAD (239 high, 239 low), and PRAD (246 high, 246 low). This analysis was conducted for all 26 hub genes; however, the total number of patients for KM analysis varied slightly for each gene across SCTs. Combining patients from all four cancer types, the approximate number of patients for each gene was around 1150 (±5) for both the high-expression and low-expression groups, depending on gene-specific factors such as data availability, expression thresholds, cutoff-based categorization, and filtering criteria in the TCGA dataset. Additionally, we used TNMplot (https://tnmplot.com/analysis/) [33] for gene expression analysis of our hub gene set to further validate the genes in a larger patient cohort. The TNMplot includes 56,938 unique samples from GEO, GTEx, TCGA, and TARGET databases (As of 17-01-2025), with 15,648 normal, 40,442 tumors, and 848 metastatic tumor samples. This large cohort provides a comprehensive view of gene expression and enhances the generalizability of our findings. We analyzed gene expression patterns in normal, tumor, and metastatic tissues for SCTs (metastatic data not available for LUAD), aiming to identify genes with significant expression changes related to metastasis and their potential roles in cancer progression.

### 2.10 Integrative Drug-Target Interaction Analysis (IDTIA)

In this study, we integrated four databases for IDTIA: DGIdb (v.5.0) [34], GETdb-2025 [35], PharmKB-2025 [36,37], and DTC-2025 [38]. These databases provide drug-target interactions, enhancing coverage and accuracy. DGIdb offers a drug-gene catalog, GETdb curates drug-target relationships, PharmKB provides pharmacogenomic details, and DTC includes clinical data. This multi-source approach allows cross-validation and increases the sensitivity of our analysis. This approach allows us to explore more drug-target interactions and advance drug discovery. The identified 26 hub genes, along with any drugs interacting with these genes, could potentially be repurposed for SCTs and offer new therapeutic options.

## 3. Results

### 3.1 Screening of Differentially Expressed Genes (DEGs)

The RNA expression data were extracted from the TCGA database to compare DEGs between “Tumor”, and “Normal” samples across SCTs. All DEGs were identified using criteria of absolute fold changes ≥ 1.5 and adjusted p-value < 0.05. The numbers of upregulated (U) and downregulated (D) genes in each cancer type were as follows: BRCA (U = 1161, D = 2479), LUAD (U = 1635, D = 2453), COAD (U = 1548, D = 2827), PRAD (U = 620, D = 973). We proceeded by selecting a set of 199 common genes (U=41, D=158) identified as significant by both the Limma and Elastic Net models, which showed similar expression patterns across breast, lung, and colorectal cancers. To ensure our findings’ reliability and biological relevance, we performed a two-step comparison between TCGA and independent GEO datasets of the SCTs. First, we examined differential expression (log2FC) by creating tumor-versus-normal heatmaps for both datasets. The results showed strong consistency in both directions in both datasets. This indicates that the transcriptional changes observed in TCGA are reproducible in GEO. Second, we compared the average gene expression levels between tumor and normal samples across the two datasets. We observed statistically significant positive correlations in all SCTs, suggesting a conserved expression pattern across different platforms and cohorts. Together, these findings confirm that the 199 common genes across the SCTs and their expression patterns are not artifacts of specific datasets but rather reflect robust and biologically consistent signals, supporting the strength and reliability of our findings. Complete details are given in **Suppl. Data-1**.

### 3.2. Gene set enrichment analysis (GSEA)

We first performed GSEA to identify common pathways among the 199 shared genes involved in key cancer hallmarks such as uncontrolled cell division, DNA damage, immune evasion, and disrupted cell communication. These pathways are consistently altered across SCTs, suggesting that these genes may contribute to tumorigenesis and could serve as potential biomarkers or therapeutic targets. Next, GSEA was applied to the entire gene expression dataset of SCTs individually to gain a broader understanding of the underlying biological processes and to identify global trends in the data. The enrichment results revealed that global gene sets are significantly associated with processes like the cell cycle, mitotic programmed cell death, chromosome maintenance, translation, transcription, DNA damage checkpoints, DNA repair, transport, signaling, and more. Additionally, we identified several common pathways altered by DEGs across SCTs, including the cell cycle, DNA repair, extracellular matrix organization, cellular responses to stress, MHC class-II antigen presentation, and membrane trafficking. These pathways were similarly affected in all cancers. Detailed pathway information is shown in **Fig. 2**.

**Figure 2:**
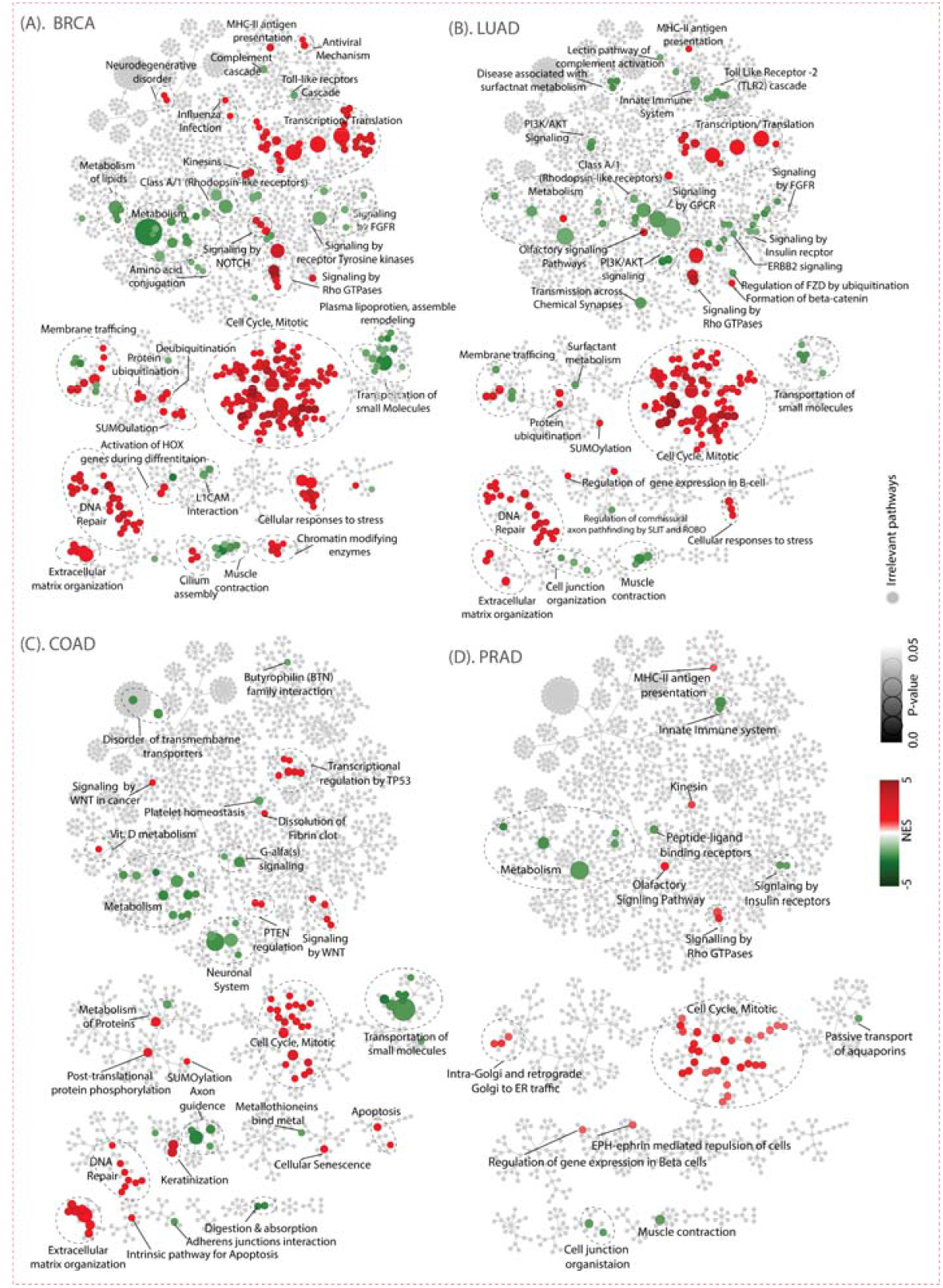
Gene set enrichment analysis (GSEA) mapped onto Reactome pathways, detailing the functional annotation of all differentially expressed genes from four cancer types. The colour of the nodes represents the normalized enrichment score (NES); positive NES values indicate increased pathway activity, while negative NES values indicate decreased activity. Each node represents a pathway. The significance of each pathway is highlighted by the node border, with darker colours indicating greater significance. Grey nodes indicate irrelevant pathways.

### 3.3. Co-expression modules detection by WGCNA

All DEGs from SCTs were analyzed using the WGCNA package in R. Soft threshold values for the datasets were chosen with a cutoff *R*^2^ value of 0.8, selecting a power value ensures that the network has the desired scale-free topology (power value ≥ *R*^2^). Module-trait relationships showed positive or negative correlations with tumors, with a high correlation between gene significance (GS) and module membership (MM), suggesting a strong association between key genes within the modules and the trait [39]. All modules across SCTs significantly correlated with the tumor group except for PRAD (Module-1-corr. ≤ 0.5). All details about co-expressed modules are depicted in **Fig. 3A**.

**Figure 3:**
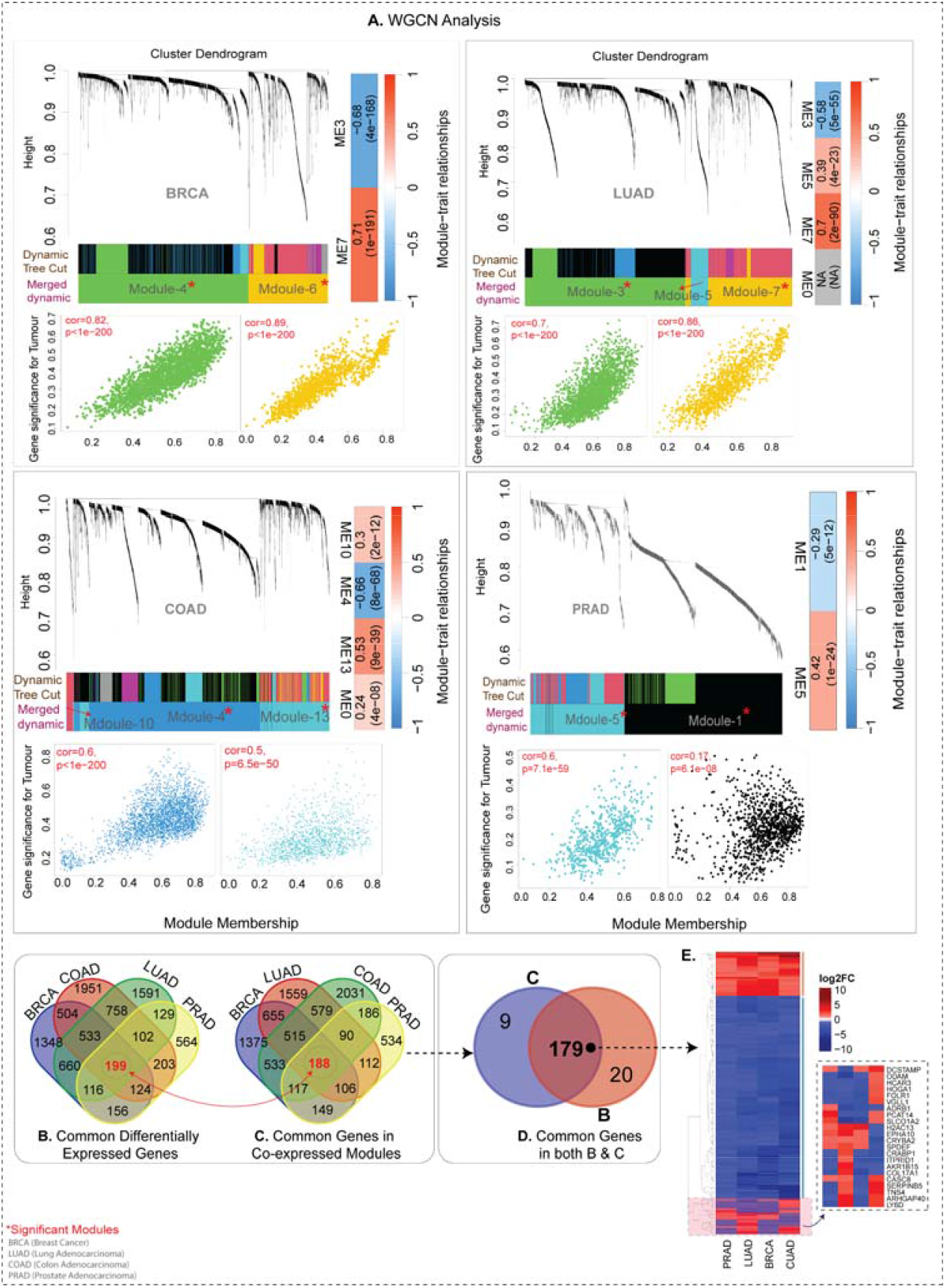
**(A)** Hierarchical clustering of genes based on the 1-TOM matrix arranges the co-expression network modules in a dendrogram for four types of cancer data (BRCA, LUAD, COAD, and PRAD). Each module is represented by a different colour. Scatterplots of gene significance (GS) for the tumor group versus module membership (MM) in the modules show a very significant correlation. This implies that the hub genes within these modules are also highly correlated with the incidence of the tumor group in the four types of cancer data. **(B-C)** Venn diagrams showing the common genes among the cancers as DEGs based on LIMMA (B) and WGCNA (C). **(D)** Common genes among the modules of all cancer types based on the combined LIMMA and WGCNA. **(E)** Heatmap showing the expression pattern of the 179 genes common across all cancer types.

Our analysis led to the identification of 199 DEGs that were common to all SCTs. We also found significant modules with a strong correlation (±0.5) to the phenotype (Tumor vs. Normal) and a p-value ≤0.05 across SCTs. Upon comparing genes across SCT modules, we identified 188 common genes, of which 179 overlapped with the 199 common DEGs obtained from LIMMA across cancers **(Fig. 3B-D)**. We cross-checked the expression patterns (log2FC) of 179 common genes across SCTs, with a heatmap showing consistency except for 22 genes **(Fig. 3E)**. These 179 genes common to SCTs and co-expressed modules suggest a crucial role in cancer progression and shared regulatory mechanisms. Detailed information about each module and their key genes, along with MM and GS scores, is provided in the **Suppl. Data-2**.

### 3.4 Protein-Protein Interaction (PPI) Network and MCL Clustering Analysis

We built a PPI network of 179 genes using BioGrid and HINT databases (considering only physical and binary interactions), comprising 2872 genes and 66,399 interactions. We calculated the topological properties (Degree, Betweenness, Closeness, and Neighbourhood connectivity) of the network to identify hub genes, located at the center of the network, which play key roles in maintaining the network’s structure and function and may be crucial for understanding biological processes or diseases. The node-degree distribution suggests that the network follows a scale-free topology [40], a common feature of biological networks **(Fig. 4A-B)**. Several genes from our list, such as *KIF20A, CAV1, OTX1, CMTM5, MYBL2, SCN2B, CTNNA3, PLP1, CRYAB, DES, FHL1, NRG1, HJURP, TPX2* etc. are among these central hub genes.

**Figure 4:**
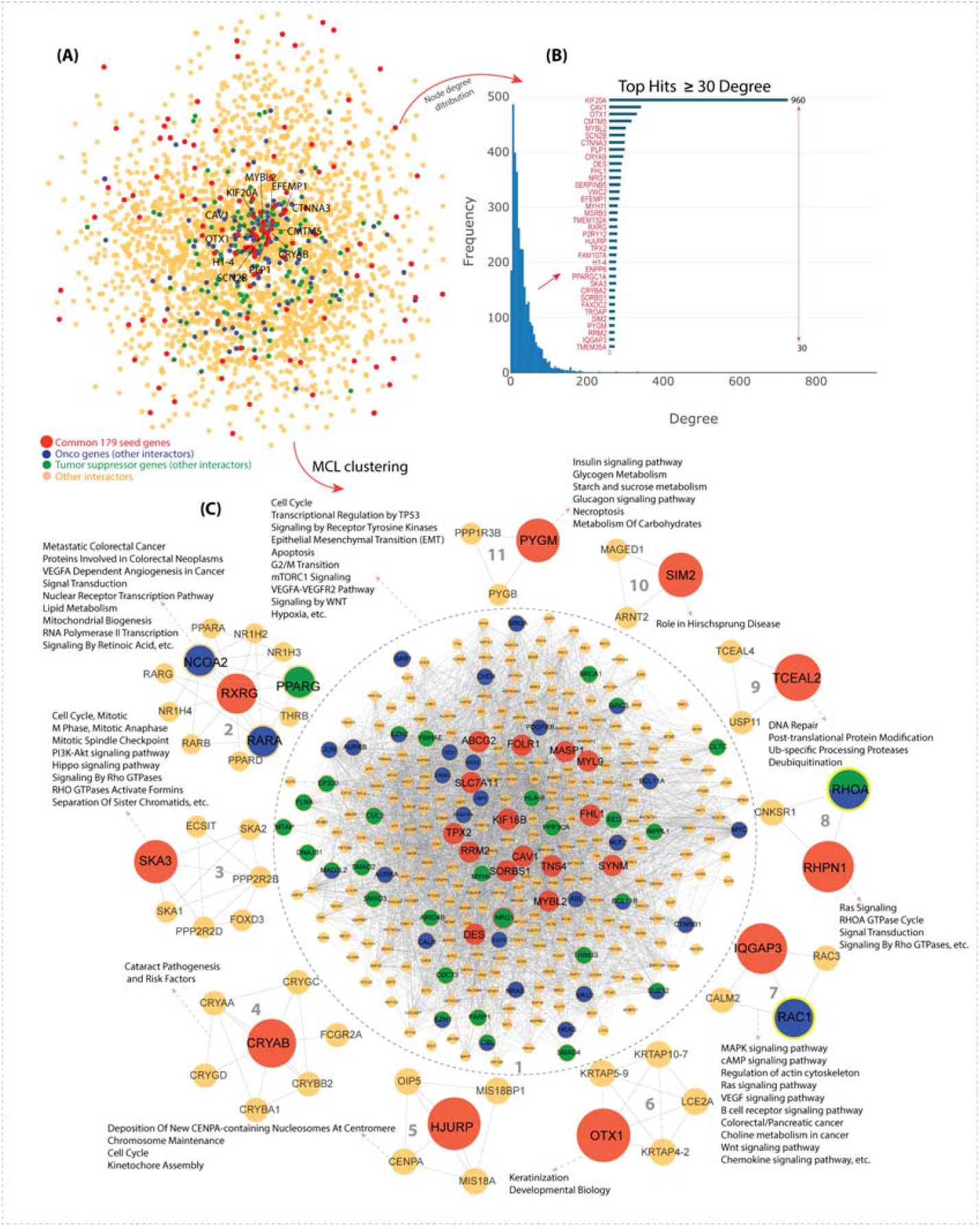
**A) PPI Network:** The Protein-Protein Interaction (PPI) network is constructed using 179 common genes from all cancer types, represented as red nodes. This network also includes their first-degree partners (orange nodes), including oncogenes (blue) and tumor suppressor genes (green). All nodes are interconnected by edges, with hub genes positioned at the centre of the network. **(B) Bar Plot Analysis:** The bar graph illustrates the node degree distribution, confirming the scale-free nature of the network, as evidenced by a few genes having high degrees while many have lower degrees. Another bar plot highlights genes with a degree ≥ 30, identifying them as hub genes within the set of 179 common genes. **(C) Functional Clustering:** The PPI network is divided into 11 functional clusters using the Markov clustering algorithm. These clusters are associated with distinct biological pathways.

We identified 11 functional clusters containing at least one hub gene from our list, clusters that do not have any hub gene were excluded from this study **(Fig. 4C)**. These functional clusters allow for a more detailed exploration of complex biological systems, providing deeper insight into how genes work together to regulate cellular processes and pathways. Dysregulation of these pathways can contribute to the development of complex diseases, including cancers. Furthermore, we conducted a pathways’ overrepresentation analysis for each cluster and found that most clusters were prominently involved in cancer-related pathways, including ‘Cell Cycle and Mitotic Regulation’, ‘Signaling Pathways’, ‘Angiogenesis’, ‘DNA Repair and Apoptosis’, and ‘Metabolic Pathways’. Additionally, the infrared genes interacting with these hub genes play significant roles in multiple cancers, highlighting the need to further investigate their functions. The network topological properties and cluster pathways overrepresentation analysis are given in **Suppl. Data-3**.

### 3.5 Immune-correlation analysis based on key genes

We analyzed Pearson correlations between 26 hub genes and immune-related genes (including immune checkpoints, chemokines, chemokine receptors, and immune stimulators/inhibitors) across SCTs. The hub genes showed positive correlation with many chemokines across SCTs, suggesting their involvement in regulating the intratumoral chemokine microenvironment responsible for immune cell recruitment, inflammation, and metastasis (**Fig. 5A**). The hub genes also showed significant correlations with two chemokine receptors (CCR10, CXCR4) in SCT, indicating their involvement in signaling pathways mediated by these receptors **(Fig. 5B)**. We further analyzed the correlation between the hub genes and immune inhibitory proteins, finding a strong positive correlation with LAG3 in SCTs. LAG3, a key checkpoint inhibitor, plays a critical role in the immunosuppressive tumor microenvironment. Other proteins, including CD160, ADORA2A, and IL10RB, showed significant negative correlations, highlighting their role in modulating the tumor microenvironment **(Fig. 5C)**. Besides, we also found a positive correlation between hub genes and immune stimulatory proteins, including TNFRSF4, IL6, MICB, TNFRSF8, CD70, CXCR4, and CD40 **(Fig. 5D)**. Additionally, we found a significant correlation between the hub genes and tumor-related Stromal_Score, Immune_Score, and ESTIMATE_Score [41,42] (**Fig. 5E**). These findings highlight the crucial role of the 26-hub genes in modulating the tumor microenvironment, potentially influencing tumor progression, immune evasion, and therapy response.

**Figure 5:**
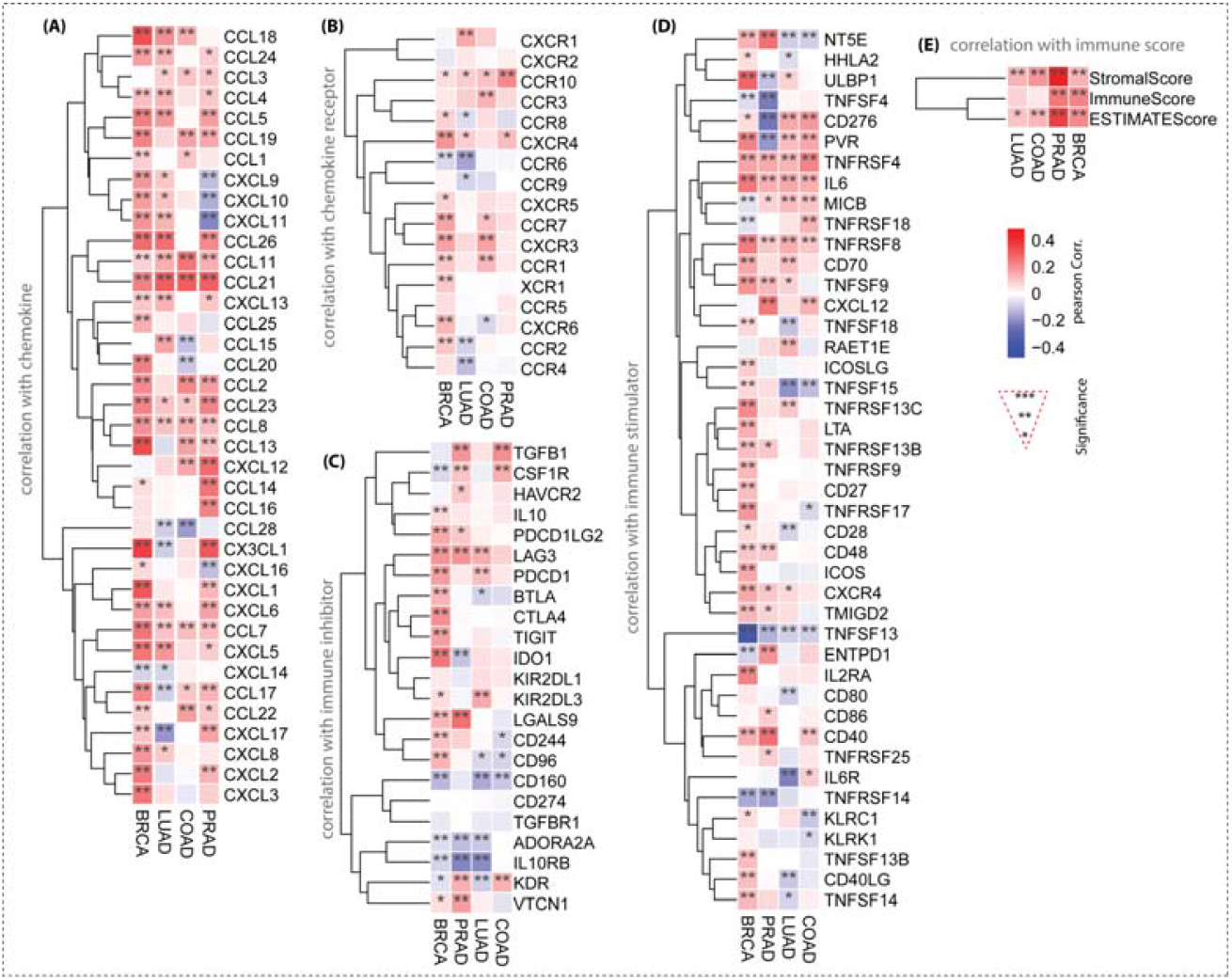
Correlation analysis between the gene set and immune-related genes: **(A)** with chemokines, **(B)** with chemokine receptors, **(C)** with immune inhibitors, **(D)** with immune stimulators, and **(E)** with the overall immune score. Significant positive correlations are shown in red, and negative correlations in blue. A high level of significance is indicated by asterisks(*).

### 3.6. Survival Analysis

The prognostic significance of the 26-hub genes in SCTs was evaluated using the GEPIA 2.0 tool. Kaplan-Meier analysis and log-rank tests revealed that many key genes are linked to poorer survival outcomes when their expression levels are high. A hazard ratio (HR) greater than 1 indicates that higher HR values correspond to an increased risk of death. Genes such as HJURP, IOGAP3, KIF18B, MYBL2, RRM2, SLC7A11, SKA3, TPX2, and FOLR1 exhibit high HR values (ranging from 2.6 to 4.5) with low p-values, signifying that their elevated expression is significantly associated with worse survival outcomes. The log-rank p-values further validate the statistical significance of survival differences between high and low-expression groups for most of these genes. These genes may serve as prognostic biomarkers, with their overexpression potentially indicating more aggressive cancer progression. Conversely, genes like ABCG2, PYGM, TCEAL2, SORBS1, DES, CRYAB, SYNM, FHL1, MASP1, SIM2, and MYL9 have HR values below 1 (ranging from 0.21 to 0.69), indicating that reduced expression of these genes correlates with poorer survival. The log-rank p-values also confirm the statistical significance of these survival differences. Lastly, Genes RHPN1, CAV1, NRG1, OTX1, TNS4, and RXRG show no significant survival differences between high and low-expression groups, as shown in the survival plot in **Fig. 6**.

**Figure 6:**
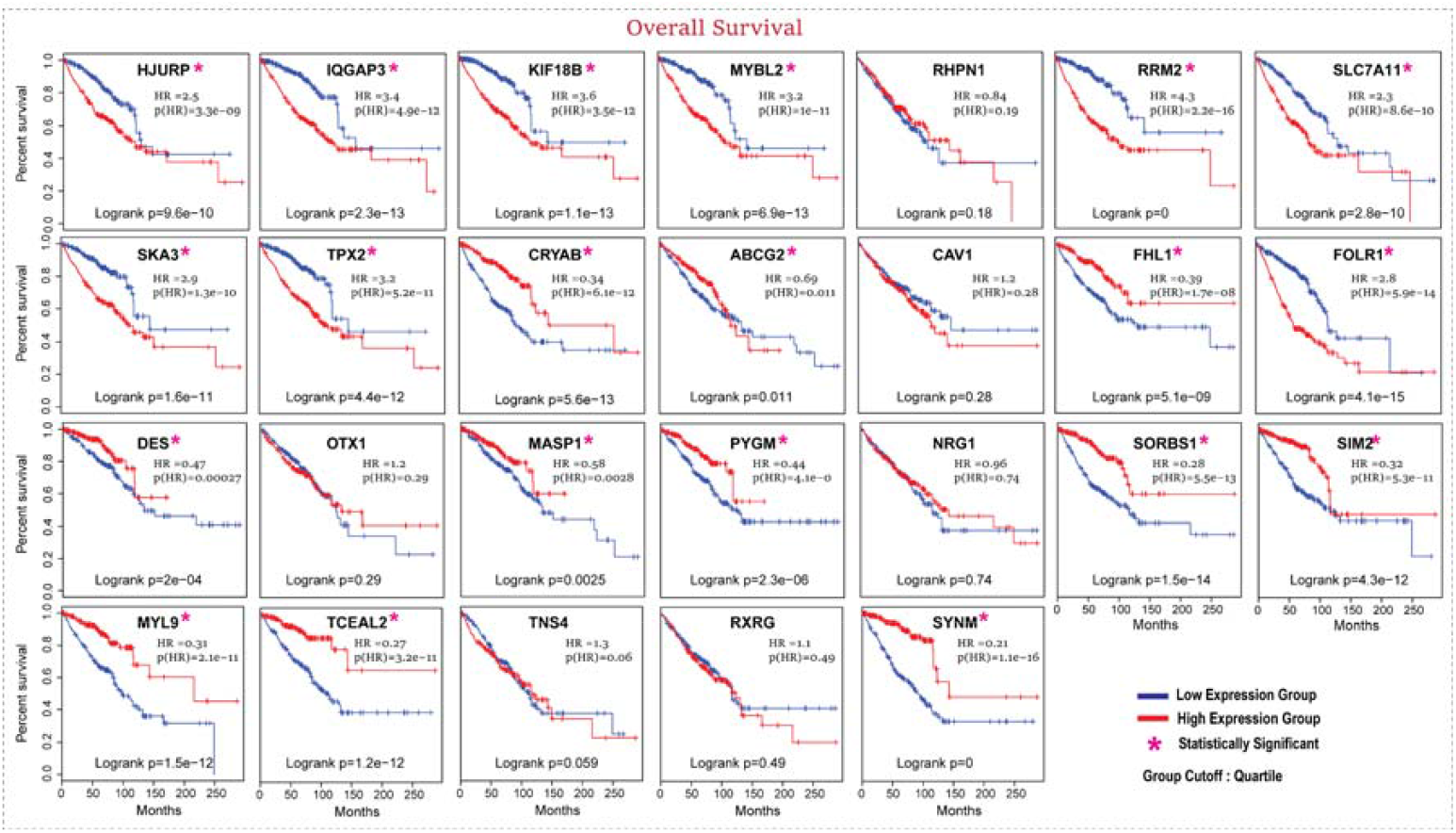
Kaplan–Meier survival curves for the 26 key genes characterizing overall survival (OS) differences in SCT, with statistical significance indicated by asterisks. The high and low expression groups, determined by quartile cutoff, are represented by red and blue lines, respectively.

### 3.7. Validation of Key Gene Expression Patterns Across a Large Cohort

TNMplot analysis showed that the expression patterns of 26-hub genes (except for SIM2) in the TCGA dataset align closely with those observed in a much larger cohort of patients, integrating data from GEO, GTEx, TCGA, and TARGET repositories. This consistency strengthens the reliability of our findings, suggesting that the identified gene expression patterns are consistent across a broader, more diverse population in SCTs. Overall, these findings underscore the potential of these 26 hub genes as valuable candidates for further investigation and as reliable potential biomarkers for diagnostic and prognostic applications. Additionally, among hub genes, we identified nine (KIF18B, RRM2, MYBL2, IQGAP3, TPX2, SLC7A11, RHPN1, HJURP, and SKA3) that show higher expression in metastatic tumors compared to primary tumors across colon, breast, and prostate cancers (**Suppl. Fig. 4**). These genes participate in critical cellular processes that directly or indirectly contribute to cancer metastasis such as E2F targets and cell cycle regulation (KIF18B, RRM2, MYBL2, TPX2, and HJURP), the RHOA GTPase cycle (RHPN1, IQGAP3), mTORC1 signaling (RRM2, SLC7A11), generic transcription pathways (TPX2, RRM2, MYBL2), and signal transduction (RHPN1, IQGAP3). These genes may provide valuable insights into tumor progression and could serve as potential biomarkers or therapeutic targets.

### 3.8. Drug-Target Interaction Analysis

Next, we studied potential interactions between genes in our key gene set and multiple classes of drugs. These interactions were analyzed by integrating four different databases, as shown in the schematic diagram (**Fig. 7A**). We identified 10 genes (ABCG2, CAV1, DES, FHL1, FOLR1, NRG1, PYGM, RRM2, RXRG, and SLC7A11) that interact with multiple FDA-approved drugs, but FHL1 was later excluded due to having only a single interaction (**Fig. 7B**). The approved drugs are currently in use for a range of diseases, including cancer (antineoplastic/chemotherapeutic agents), infections (antibiotics and antimicrobials), hormone-related conditions (hormonal medications and endocrine agents), and inflammatory disorders (anti-inflammatory and related agents). These genes also interact with other drugs, including natural compounds and experimental or investigable drugs **(details are given in Suppl. Data-4 and Suppl. Fig. 5)**. For this analysis, we focused on FDA-approved drugs and natural compounds because these substances have already been tested for safety and efficacy, and are more immediately applicable for therapeutic use. As FDA-approved drugs are established for clinical use, they can be readily repurposed for new treatments. Natural compounds have a long history of medicinal use and are considered safe for human consumption. Focusing on these compounds gives a more direct path to clinical application and reduces the risks associated with experimental or investigable drugs that have not been fully validated. Two of the genes (‘PYGM’ and ‘ABCG2’) in the identified key gene set were also found to interact directly with several classes of natural compounds **(Fig. 7C)** that show promising potential as antineoplastic/chemotherapeutic agents, steroid/hormone derivatives, monoclonal antibodies/antibody-drug conjugates, enzyme inhibitors, growth factors/ modulators and antiviral agents. The overall drug interaction details are shown in **Fig. 7D-E**.

**Figure 7:**
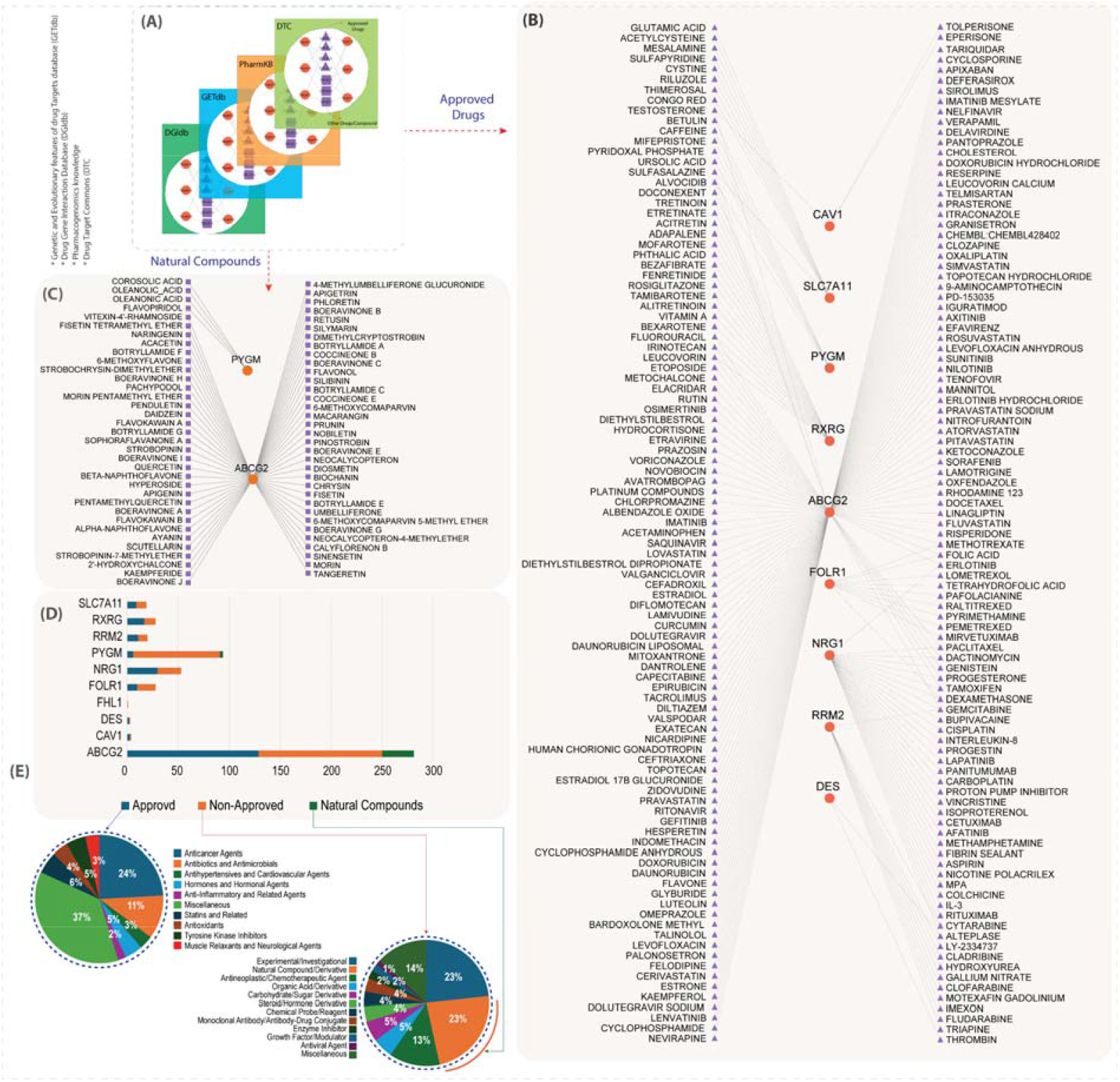
Integrative Drug-Target Interaction Analysis (IDTIA): **(A)** Schematic representation of the integration of four databases to get the Drug-Target relationship. **(B)** The list of FDA-approved drugs that are directly involved in 10 key target genes. **(C)** The list of natural compounds that interacted with two target genes namely ‘PYGM’ and’ABCG2’. **(D)** A bar graph showing the distribution of drug interaction with target genes. **(E)** Pie charts showing the categories of approved drugs (upper left chart) and non-approved drugs/natural compounds (lower right chart).

## 4. Discussion

Cancer is a major global health problem, with breast, lung, colon, and prostate cancers being the most commonly diagnosed with associated high mortality. Each cancer type has unique molecular signatures, making it difficult to use the same biomarkers across different cancers. A common set of biomarkers could streamline diagnosis, enabling faster and easier cancer detection and classification. These common potential biomarkers could aid in developing therapies effective across multiple cancers, minimizing the need for cancer-specific treatments and accelerating the development of new therapies.

In this study, we analyzed TCGA datasets to identify common DEGs across SCTs, revealing that most of the DEGs are involved in cell cycle, translation, transcription, DNA damage checkpoints, and DNA repair. These processes play a crucial role in uncontrolled cell division, altered protein synthesis, and bypassing DNA damage checkpoints, well-known hallmarks of cancer in cancer biology [43,44]. Targeting these pathways could effectively disrupt cancer growth and proliferation across different types.

We performed WGCNA to identify correlated gene modules within SCTs and found 188 common genes across each cancer module. By comparing DEGs from the LIMMA pipeline with WGCNA results, we identified 179 common DEGs that are highly co-expressed and likely crucial for cancer progression. Using protein-protein interaction networks, we identified 11 functional clusters and 26 hub genes, including *‘RRM2’, ‘MYBL2’, ‘TPX2’, ‘DES’, ‘KIF18B’, ‘SYNM’, ‘SLC7A11’, ‘TNS4’, ‘FOLR1’, ‘ABCG2’, ‘SORBS1’, ‘MASP1’, ‘CAV1’, ‘MYL9’, ‘FHL1’, ‘RXRG’, ‘SKA3’, ‘CRYAB’, ‘HJURP’, ‘OTX1’, ‘IQGAP3’, ‘RHPN1’, ‘TCEAL2’, ‘SIM2’, ‘PYGM’* and *‘NRG1’* which have high network centrality and play a central role in regulating biological processes.

Further, we analyzed 11 distinct functional clusters within the PPI network, each representing genes and pathways associated with cancer progression. For instance, <u>Cluster-1</u>, the largest, contains 15 hub genes interacting with 286 others, including 29 oncogenes and 28 tumor suppressor genes, enriched in cancer pathways like the cell cycle [45], EMT [46], mTORC1 signaling [47], VEGFA-VEGFR2 [48,49], WNT [50], and hypoxia [51]. <u>Cluster-2</u> has 12 genes, with RXRG as the hub, linking immune regulation and inflammation in cancer, enriched in pathways like metastatic colorectal cancer, angiogenesis, lipid metabolism, and retinoic acid signaling [52–54]. <u>Cluster-3</u> focuses on SKA3, linked to cell cycle [45], PI3K-Akt [55,56], and Rho GTPase signaling [57] with high expression correlating with poor outcomes. Cluster-4 focuses on CRYAB, involved in cataract pathogenesis, apoptosis, inflammation, and oxidative stress [58], influencing tumor development through the tumor microenvironment. <u>Cluster-5</u>, with HJURP as the hub, is enriched in chromosome maintenance and centromere nucleosome deposition, with HJURP as a potential oncogene and biomarker [59,60]. <u>Cluster-6</u> highlights OTX1 and its interactors, involved in keratinization, development, and several cancers [61–64], making them potential therapeutic targets. <u>Cluster-7</u> features IQGAP3 and interactors like RAC1, RAC3, and CALM2, participating in MAPK [65], VEGF [66], and Wnt signaling [67]. <u>Cluster-8</u>, with RHPN1 as the hub, is crucial in Ras signaling [68] and the RHOA GTPase cycle [57]. <u>Cluster-9</u> involves TCEAL2 (hub gene), TCEAL4, and USP11, which are linked to DNA repair [69], protein modification [70], and Deubiquitination [71]. Cluster-10, with SIM2 as the hub, is associated with Hirschsprung disease and cancer [72], with ARNT2 promoting angiogenesis [73,74] and MAGED1 supporting apoptosis [75,76]. Finally, <u>Cluster-11</u> involves PYGM (hub gene), PYGB, and PPP1R3B, linked to cancer metabolism through insulin and glycogen pathways [77–79], where PYGM is abnormally expressed in tumors, PYGB drives proliferation in pancreatic and liver cancer [80], and PPP1R3B alters glycogen metabolism to support cancer progression [81]. <u>A comprehensive discussion of all 11 modules and their corresponding hub genes, including relevant references, is presented in</u> **Suppl. Data-5**.

In this study, we focused on 26 hub genes that were differentially expressed (DEGs), many of which are well-established in the literature for their roles in various cancer-related processes [details are given in supplementary data 5]. However, we also identified several key inferred genes—such as *UBC, ANLN, KIF14, VIRMA, ESR1, CIT, MYC, CUL3, JUN, BIRC3, PRC1, KIF23, ECT2, PARK2, TP53*, among others—that exhibit strong connectivity with these hub genes within the PPI network. Notably, in several biological pathways, some hub genes do not directly participate but influence the network indirectly through their interactions with these inferred genes. These inferred genes are implicated in a wide range of critical hallmark pathways, including “cell cycle and proliferation”, “cell signaling and growth regulation”, “cell death and stress response”, “metabolism and energy regulation”, “differentiation and development”, “hormone and receptor-mediated signaling”, “immune and inflammatory responses”, and “cell structure and interactions”. As such, it is essential to consider the functional relevance of these inferred genes, even if they are not categorized as DEGs or hub genes, because they play pivotal roles in the broader molecular network and contribute significantly to the regulation of key biological processes. The details of inferred genes are given in **Suppl. Data-3**.

The drug-target interaction analysis revealed that key genes like *ABCG2, NRG1, RXRG, PYGM*, and *RRM2* interact with multiple drugs, including FDA-approved medications, natural compounds, and experimental drugs. Many of these approved drugs are already used as anticancer agents, as well as for conditions like infections, hypertension, cardiovascular diseases, hormonal imbalances, inflammation, and neurological disorders. This includes antibiotics, antihypertensives, hormones, anti-inflammatory agents, statins, antioxidants, tyrosine kinase inhibitors, and muscle relaxants, which could serve as potential therapeutics for SCTs, with repurposing accelerating cancer treatment development by leveraging existing safety and efficacy data.

Additionally, TNMplot enabled us to analyze DEGs between primary and metastatic tumors across SCTs, except for LUAD where metastatic data was unavailable. Our analysis revealed a subset of 9 genes (*KIF18B, RRM2, MYBL2, IQGAP3, TPX2, SLC7A11, RHPN1, HJURP*, and *SKA3*) from the 26-hub genes that showed differential expression between primary and metastatic niches, all of which have been previously recognized for their importance in carcinogenesis. For instance, KIF18B is a member of the kinesin-8 proteins and is involved in spindle assembly and the separation of chromosomes during mitosis. KIF18B is upregulated in many human cancers and is associated with a poor prognosis due to its impact on resistance to chemotherapy and radiotherapy [82–84]. Similarly, TPX2 is a microtubule protein that also functions in mitotic spindle assembly. It has been identified as a potential prognostic marker in multiple cancers, including colorectal carcinoma, hepatocellular carcinoma, neuroblastoma, gastric cancer, breast cancer, and bladder cancer [85]. Furthermore, the rhophilin Rho GTPase binding protein 1 (RHPN1) plays a role in signal transduction involving Rho GTPases and is involved in cytoskeletal organization and, hence, cancer metastasis [86]. The closely related RHPN1-AS1 gene is classified as a long non-coding RNA (lncRNA) and is transcribed in the antisense orientation relative to the RHPN1 gene. It acts as an oncogene, promoting cell proliferation, migration, and invasion, particularly in cancers like lung adenocarcinoma, ovarian cancer, gastric cancer, and glioblastoma [87]. RHPN1-AS promotes carcinogenesis by sponging miRNAs, enabling oncogenic proteins like TPX2 to drive cancer cell proliferation, adhesion, and migration. SKA3, an oncogene, regulates chromosome separation and cell division and is associated with lung adenocarcinoma, as well as prostate, cervical, and breast cancers [88].

RRM2, the catalytic subunit of ribonucleotide reductase, is essential for DNA replication and repair. It is upregulated in cancers such as kidney, lung, and liver, and its expression is linked to poor prognosis [89–91]. Similarly, MYBL2 has been shown to act as an oncogene and carries a poor prognosis in multiple cancers [92]. In an ovarian cancer mouse model, MYBL2 collaborates with CCL2 chemokine to sustain the immunosuppressive tumor microenvironment and resistance to anti-PDL1 therapy by inducing the infiltration of immunosuppressive macrophages [93]. HJURP (Holliday_Junction_Recognition_Protein) is an oncogene that functions as a chaperone of histone H3, HJURP is upregulated in cancers like prostate, breast, renal, and oral cancers. It promotes tumor proliferation, invasion, and metastasis, and alters the immune landscape within tumor tissue [94]. IQGAP3, a Rho family GTPase, regulates key cancer processes like proliferation, apoptosis, migration, invasion, and angiogenesis. Its pro-tumorigenic role makes it a potential therapeutic target in various cancers [95], including high-grade serous ovarian cancer, which has the highest mortality rate in women [96]. SLC7A11, a cystine transporter, helps cancer cells produce glutathione and other compounds, protecting against oxidative stress-induced cell death[97]. It plays a role in tumorigenesis by regulating redox homeostasis, cell proliferation, immune modulation, metastasis, and therapeutic resistance [98].

Further analysis revealed that most genes in this group are co-expressed, indicating functional relationships and likely act together in a coordinated manner. Genes such as *TPX2, RRM2, HJURP*, and *MYBL2* from this group are involved in key pathways including the cell_cycle, gene_expression_regulation, Rho_GTPase_signaling (*RHPN1; IQGAP3*), mTORC1_signaling (*RRM2; SLC7A11*), and the E2F_targets pathway (*KIF18B; RRM2; MYBL2*). Dysregulation of these pathways due to altered gene expression can contribute significantly to cancer development, progression, and metastasis.

Additionally, we analyzed the expression of these nine genes across 33 TCGA-Pan cancer types and found they are significantly overexpressed in more than 18 cancers. This suggests their role as oncogenes or tumor markers, potentially serving as potential biomarkers for early detection, prognosis, treatment response, and drug targets, allowing us to better predict their involvement in tumor initiation, progression, and metastasis.

In summary, our study identifies key hub genes involved in cancer progression, with several associated with metastasis and immune regulation. These genes may serve as prognostic biomarkers and therapeutic targets, with drug-gene interaction analysis highlighting the potential for drug repurposing. Further experimental validation is needed to confirm their clinical relevance in cancer diagnosis, prognosis, and treatment.

## 5. Conclusion

In this study, we uncovered a shared molecular signature of 179 key genes across four major cancer types—breast, lung, colon, and prostate—offering a significant opportunity to streamline diagnostics and enable unified therapeutic strategies. Among these, 26 hub genes were identified as central players in PPI networks, participating in key hallmark cancer pathways and cellular processes. Notably, nine of these hub genes were consistently upregulated in metastatic tumors and were associated with immune regulation, suggesting their involvement in tumor progression and immune evasion. These genes were also overexpressed in over 18 cancer types, highlighting their fundamental role in tumorigenesis and reinforcing their potential as pan-cancer biomarkers and therapeutic targets. Furthermore, drug-target interaction analysis revealed that 10 hub genes interact with FDA-approved or investigational drugs, opening avenues for drug repurposing and the development of targeted therapies. This integrative approach not only deepens our understanding of shared molecular mechanisms in cancer but also paves the way for personalized, cross-cutting treatment strategies that may apply to other complex diseases as well. As we continue to bridge genomic insights with clinical application, these findings have the potential to simplify cancer diagnostics, enhance prognostic accuracy, and optimize therapeutic pipelines. However, further experimental validation and clinical studies are necessary to confirm the diagnostic, prognostic, and therapeutic value of these candidate genes, ultimately contributing to the realization of precision oncology.

## Acknowledgment

This work was supported by a grant from ASPIRE, the technology program management pillar of Abu Dhabi’s Advanced Technology Research Council (ATRC), via the ASPIRE Precision Medicine Research Institute Abu Dhabi (ASPIREPMRIAD) award grant #VRI-20-10 to BK A-R. Additional support was provided by Al Jalila Foundation (Dubai, UAE) and UAEU Program for Advanced Research (UPAR) grants #31R088 (to BK A-R) and #12F061 (to UK).

## Author contributions

AA conceived the study design, performed the data acquisition and analysis, curated the figures, and wrote the manuscript; MFS contributed to the study design and helped in drafting the manuscript. RH provided critical input on methodology and reviewed the manuscript. UK reviewed the manuscript; MJFC provided scientific insight and critical input and edited the manuscript. BKA-R conceived the study, acquired the funding for the study, supervised the project, and wrote the manuscript. All authors read and approved the final version.

## Data Availability

All the supporting data files (Cytoscape files, Supplementary data, R scripts) are publicly available on GitHub (https://github.com/aftab3866/Suppli_Data_2025).

