## Supplementary material for "A Systems Biology Approach to Unveil Shared Therapeutic Targets and Pathological Pathways Across Major Human Cancers": Suppl. Data-5

------------------------------------------------------------------------------------------------------------

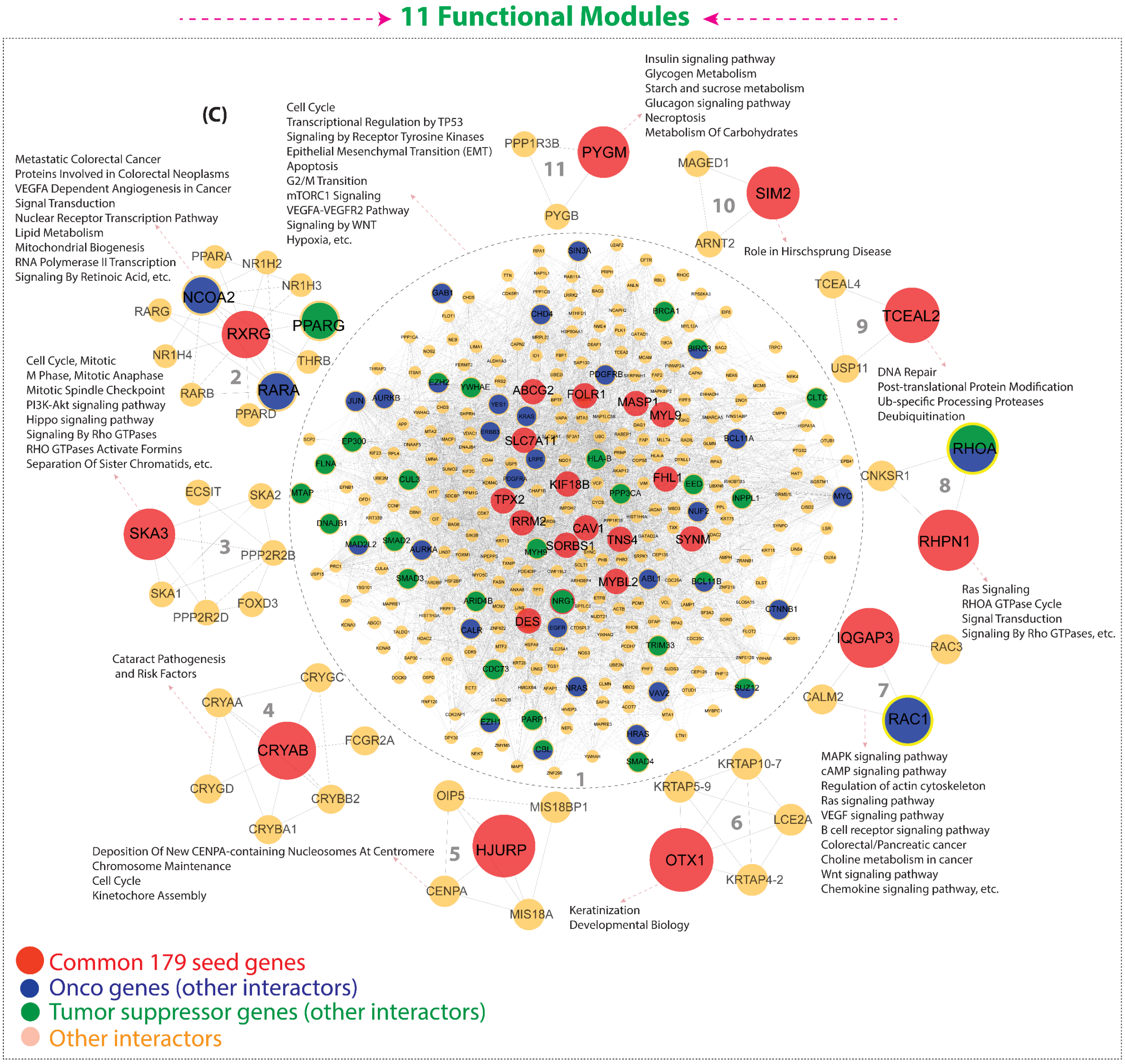


1. **Cluster-1:** This is the biggest cluster and contains 15 hub-genes ('RRM2', 'MYBL2', 'TPX2', 'DES', 'KIF18B', 'SYNM', 'SLC7A11', 'TNS4', 'FOLR1', 'ABCG2', 'SORBS1', 'MASP1', 'CAV1', 'MYL9', 'FHL1') that directly interact with other 286 genes including 29 oncogenes (OG) and 28 tumor suppressor genes (TSG) with 7 genes having a dual role as both OG and TSG, including

‘CBL’, ‘EZH1’, ‘BCL11B’, ‘SUZ12’, ‘MAD2L2’, ‘EZH2’ and ‘BIRC3’ dependent on the cellular context, such as mutation type, tissue type, or signaling environment.

In this cluster, high-degree nodes (genes) are ‘UBC’, ‘MYC*’, ‘JUN*’, ‘CIT’, ‘CAV1*’, ‘BIRC3*’, ‘HSPA8’, ‘CUL3’, ‘KIF23*’, ‘ANLN*’, ‘PRC1*’, ‘RBBP4*’, ‘MCM2*’ and ‘MYBL2*’. These genes are highly (*) or likely linked to cancers, either functioning as oncogenes, tumor suppressors, or crucial components in cellular processes, which when disrupted, may drive cancer progression. Moreover, we checked the over-representation of the gene set from this cluster in various pathways and we found that genes from this cluster are directly involved in cancer-related pathways such as ‘Cell Cycle’/‘G2/M Transition’ [1], ‘Transcriptional Regulation by TP53’ [2], ‘Signaling by Receptor Tyrosine Kinases’ [3], ‘Epithelial to Mesenchymal Transition (EMT)’ [4], ‘Apoptosis’ [5], ‘mTORC1 Signaling’ [6], ‘VEGFA-VEGFR2 Pathway’ [7], [8], ‘Signaling by WNT [9]’, ‘Hypoxia [10]’, and others. Notably, in all these cancer-related pathways, our identified hub genes are directly involved.

2. **Cluster-2:** This cluster comprises 12 genes, including two oncogenes (RARA and NCOA2) and one tumor suppressor gene (PPARG). All these genes interact with the hub gene, RXRG, which itself plays a role in tumor suppression [11]). Retinoid X Receptor Gamma (RXRG) and other partners in this cluster are associated with nuclear receptor (NR) superfamily, which play a key role in the tumor microenvironment (TME) through regulation of immune responses and inflammation [12]. This cluster is enriched in key cancer pathways such as ‘Metastatic Colorectal Cancer’, 'Involved in Colorectal Neoplasms', 'VEGFA Dependent Angiogenesis in Cancer', 'Signal Transduction' [13], 'Nuclear Receptor Transcription Pathway' [14], 'Lipid Metabolism' [15], [16], 'Mitochondrial Biogenesis'[17], 'RNA Polymerase II Transcription'[18], 'Signaling By Retinoic Acid' [19], and ‘Post-translational Protein Modification’ [20], [21].

3. **Cluster-3:** This cluster contains only one hub gene (SKA3) that interacts with four other genes ('PPP2R2D’, ‘SKA2’, ‘SKA1’, and ‘PPP2R2B’). SKA protein family members are involved in tumorigenesis and cancer progression [22], [23]. SKA3 is indirectly involved in many cell cycle pathways [1], PI3K-Akt signaling pathway [24], [25] and signaling by Rho GTPases [26]. High expression of SKA3 is linked to worse outcomes in cancer, and could be a promising target for new treatments [27].

4. **Cluster-4:** This cluster also contains only one hub gene (CRYAB) that interacts with six other genes (‘CRYBB2’, ‘FCGR2A’, ‘CRYGC’, ‘CRYAA’, ‘CRYGD’ and ‘CRYBA1). This cluster is entirely involved in ‘Cataract Pathogenesis and Risk Factors’. It has been suggested that early-onset cataracts are associated with insufficient antioxidative activity, and, therefore, a potential risk of cancer [28]. The CRYAB is a member of the small heat shock protein family, CRYAB proteins are involved in a range of signaling pathways including apoptosis, inflammation, and oxidative stress [29], and also modulate tumor development *via* the TME [30]. Therefore, CRYAB could be a potential target for cancer therapy.

5. **Cluster-5:** This cluster contains five genes that interact with one hub-gene (HJURP). This cluster is enriched in pathways such as 'Deposition of New CENPA-containing Nucleosomes at Centromere', 'Chromosome Maintenance [31]', 'Cell Cycle [1]' and 'Kinetochore Assembly [32]'. Holliday junction protein (HJURP) may act as an oncogenic factor. Its expression and immune infiltration characteristics could serve as valuable biomarkers for cancer detection, prognosis, and

treatment [33], [34]. In this cluster, we should also focus on the interactors of HJURP—such as ‘CENPA’, ‘MIS18BP1’, ‘MIS18A’, and ‘OIP5’—in tumorigenesis and their effects. This could provide new insights into targeting HJURP as a promising cancer treatment strategy.

6. **Cluster-6:** This cluster is a 6-member group with a single hub gene (‘OTX1: Orthodenticle homeobox 1’) that interacts with other genes, including 'KRTAP10-7', 'KRTAP4-2', 'KRTAP5-9', and 'LCE2A'. The genes of this cluster are involved in 'Keratinization [35], [36]' and 'Developmental Biology [37]' pathways. However, the hub gene (‘OTX1’) in this cluster is well known for its role in various human cancers [38], [39], [40], [41]. Therefore, OTX1 could be a potential target for cancer therapy; however, other interactors of OTX1 are also important because they probably contribute to the regulation of cancer-related functions of OTX1. Understanding their roles can offer a deeper insight into how this cluster functions in cancer, uncovering additional therapeutic targets for cancer.

7. **Cluster-7:** This is also a significant four-member cluster consisting of ‘IQGAP3’ (hub gene), ‘RAC1’, ‘RAC3’, and ‘CALM2’. This cluster is mostly involved in cancer-related pathways including ‘MAPK signaling pathway’ [42], ‘cAMP signaling pathway’ [43], ‘Regulation of actin cytoskeleton’ [44], ‘Ras signaling pathway’ [45], ‘VEGF signaling pathway’ [46], ‘B cell receptor signaling pathway’ [47], ‘Colorectal/Pancreatic cancer’, ‘Choline metabolism in cancer’, ‘Wnt signaling pathway’ [48], and ‘Chemokine signaling pathway’ [49]. IQGAP3 is involved in cancer proliferation, metastasis, and serves as a potential prognostic marker. It also plays critical roles in tumor progression and immune responses across various cancer types [50], [51], [52], [53]. IQGAP3 can have dual properties, acting as a pro-tumorigenic gene as well as an anti-tumorigenic gene, depending on the context. Collectively, these findings suggest that ‘IQGAP3’ is a promising therapeutic target for cancer treatment or intervention. Moreover, the interaction partners could also be important, such as ‘RAC1’ which is considered as a potential target for the prevention and treatment of cancer [54], [55], [56]. Similarly, ‘RAC3’ functions as an oncogene that promotes cancer cell proliferation and invasion [57]. Calmodulin 2 (CALM2) is a calcium-binding protein linked to the development of several cancers [58]. The CALM2 gene influences the JAK2/STAT3/HIF-1/VEGFA pathway and supports macrophage activity by modulating inflammation, angiogenesis, and macrophage polarization, which helps metastasis and promotes angiogenesis [59]. Therefore, this cluster is highly significant and contains four potential therapeutic targets for cancer.

8. **Cluster-8:** This 3-member cluster, consisting of ‘RHPN1’ (hub gene), ‘RHOA’, and ‘CNKSR1’, is also highly significant because of their involvement in several cancer related pathways sucha as ‘Ras Signaling’[45], ‘RHOA GTPase Cycle [26]’ and ‘Signal Transduction [13]’. The closely related RHPN1-AS1 gene is classified as a long non-coding RNA and has been implicated in triggering autophagy and apoptosis in prostate cancer cells by affecting the miR-7-5p/EGFR/PI3K/AKT/mTOR signaling pathway [60]. RHPN1-AS1 promotes ovarian cancer development by acting as a sponge for miR-6884-5p, which in turn, releases TOP2A mRNA, aiding in tumor progression [61]. RHPN1-AS1 promotes endometrial cancer progression by activating the ERK/MAPK pathway [62]. RHPN1-AS1 enhances cell proliferation, invasion, and migration in cervical cancer by regulating the miR-299-3p/FGF2 axis [63]. These studies highlight this gene as a potential therapeutic target for cancer.

9. **Cluster-9:** This is also a 3-member cluster, consisting of 'TCEAL2' (hub gene), 'TCEAL4', and 'USP11'. This cluster is also involved in cancer related pathways like 'DNA Repair' [64], 'Post-translational Protein Medication' [65], 'Ub-specific Processing Proteases [66]' and 'Deubiquitination [67]'. Except for 'USP11', not much information is available about these genes with respect to their involvement in biological pathways. However, as per the 'guilt by association' approach [68], [69] other genes in this cluster may also play a role in the same pathways along with 'USP11'. For example, 'TCEAL2' (hub gene) has been shown to function as a tumor suppressor in renal cell carcinoma [70]. TCEAL2 has also been found to be significantly associated with various immune cells, particularly NK cells, as well as immune-related genes in many cancers [71]. Therefore, TCEAL2 may serve as a novel prognostic marker and could influence tumor behavior through regulating immune cell infiltration.

10. **Cluster-10:** This cluster also consists of three members: 'SIM2' (the hub gene), 'ARNT2,' and 'MAGED1.' This cluster is mainly involved in Hirschsprung disease-relate pathways. There is evidence for a connection between Hirschsprung disease (HSCR) and cancer, potentially through genetic and developmental pathways [72]. HSCR has been found to be associated with mutations in several genes, including RET. Mutations in the RET gene are linked to multiple endocrine neoplasia type 2 (MEN2), a syndrome that increases the risk of medullary thyroid carcinoma and other tumors [73], [74]. At the gene level, 'SIM2' is a transcription factor and has a role in various cancers like prostate [75]. It also has been seen that 'SIM2' is overexpressed in endometrial carcinoma, with its DNA methylation and CNV (copy number variation) alterations linked to immune cell infiltration and prognosis. SIM2 is involved in transcription regulation and signal transduction, promoting cell proliferation, migration, and invasion [76]. There is a cross-talk between SIM2s and NFκB plays a regulatory role in the expression of cyclooxygenase 2 (COX-2) in Breast cancer [77]. Another gene from this cluster, 'ARNT2', which is involved in carcinogenesis, cancer progression and tumor angiogenesis [78], [79], [80]. Additionally, 'MAGED1', also from this cluster, is a member of the melanoma antigen gene (MAGE) family. Most of the genes of this family encode tumor specific antigens. 'MAGED1' contributes to anti-tumorigenesis and apoptosis across various cell types. It's down-regulation has been observed in tumor cells, including those in breast carcinoma, glioma stem cells, and colorectal cancer tissues [81], [82]. Thus, the genes in this cluster seem highly significant and could serve as both prognostic markers and therapeutic targets for various cancers.

11. **Cluster-11:** This is another 3-member cluster: 'PYGM' (the hub gene), 'PYGB,' and 'PPP1R3B.' This cluster is predominantly associated with cancer-related pathways such as 'Insulin signaling pathway' [83], [84], 'Glycogen metabolism' [85], 'Starch and sucrose metabolism [86]', 'Glucagon signaling pathway' [87], 'Necroptosis [88]' and 'metabolism of carbohydrates' [89]. However, PYGM's abnormal expression itself is associated with a variety of tumors [90]. Another gene of this cluster 'PYGB,' in involved in pancreatic cancer (PC) [91], hepatocellular carcinoma (HCC) [90]. 'PYGB' enhances cell proliferation, invasion, and metastasis, resulting in poor patient prognosis. Additionally, PPP1R3B is involved in glycogen synthetase (GYS) activation and inactivation of glycogen phosphorylase, resulting in abnormal glycogen metabolism that helps in cancer progression [92].
