## Supplementary figures and images for "A Systems Biology Approach to Unveil Shared Therapeutic Targets and Pathological Pathways Across Major Human Cancers"

### Suppl. Fig. 1

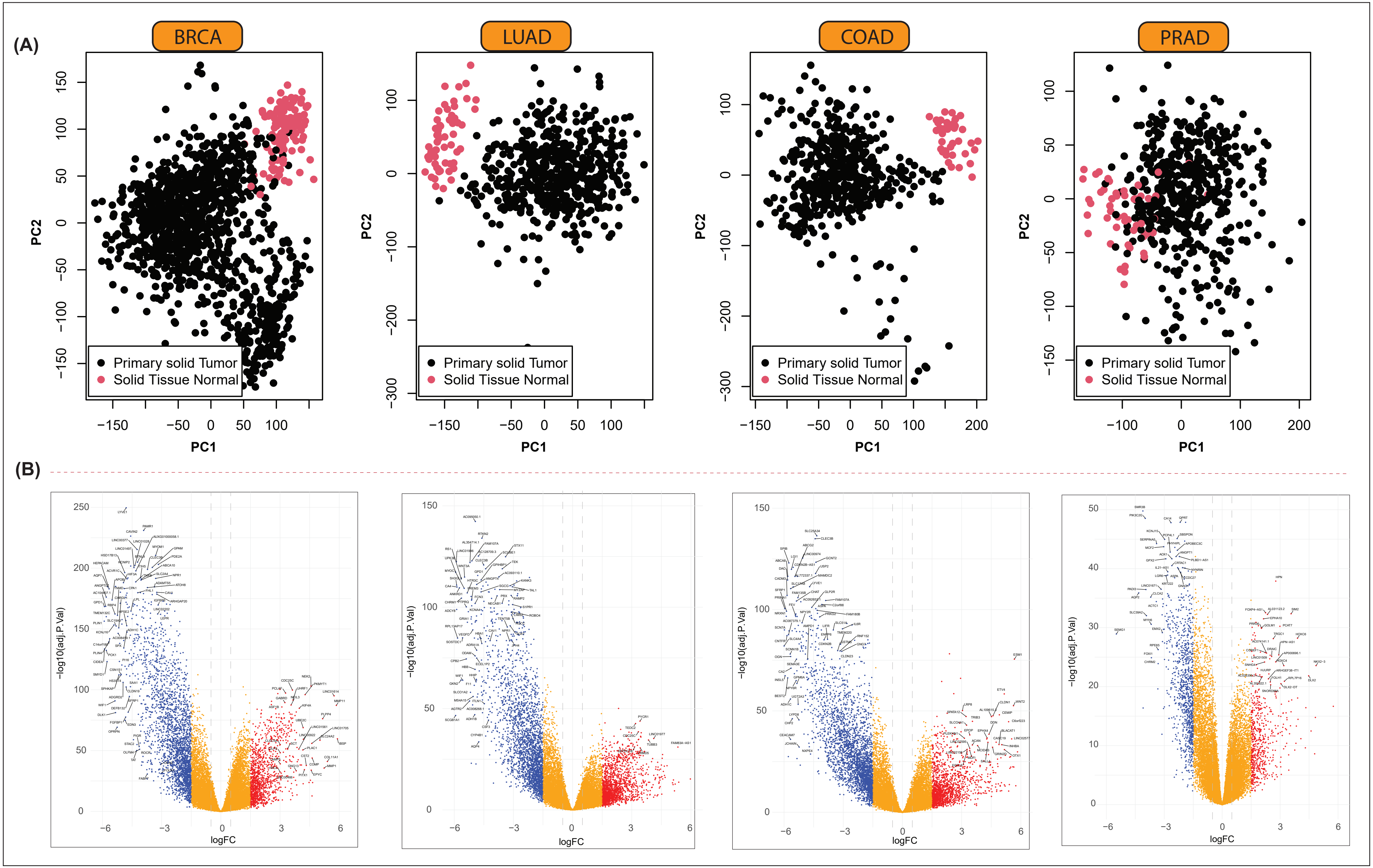

### Suppl. Fig. 2

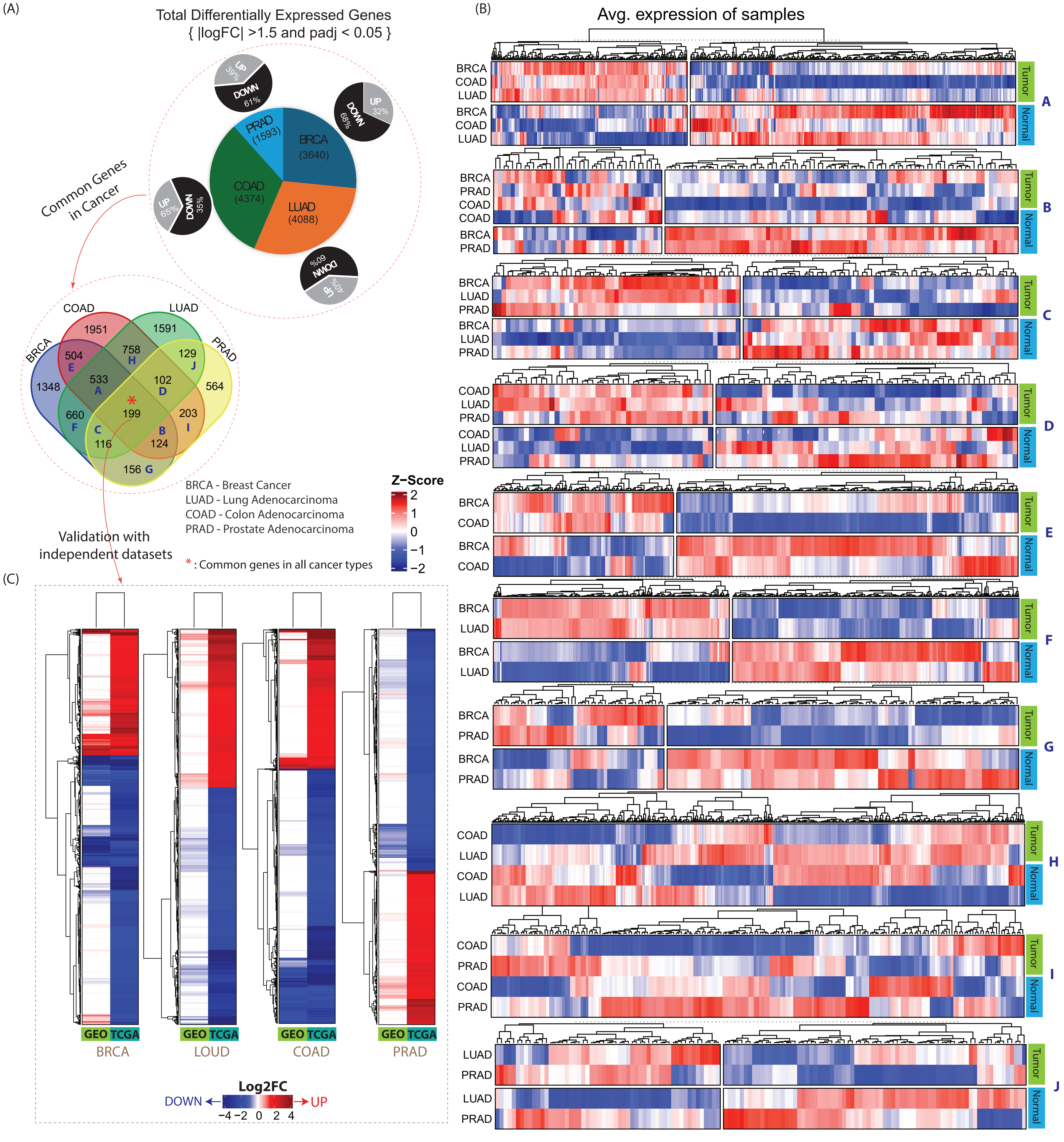

### Suppl. Fig. 4

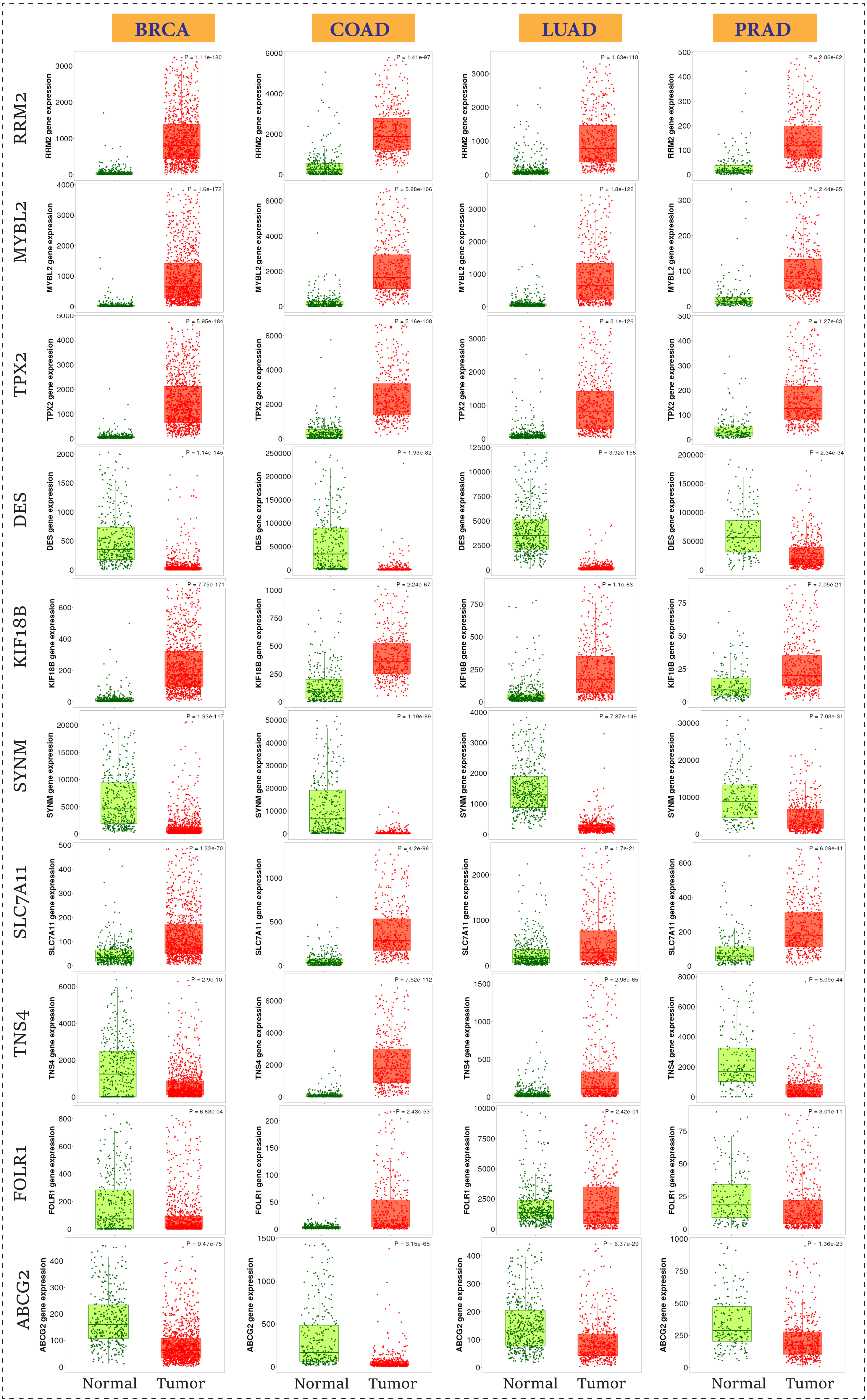

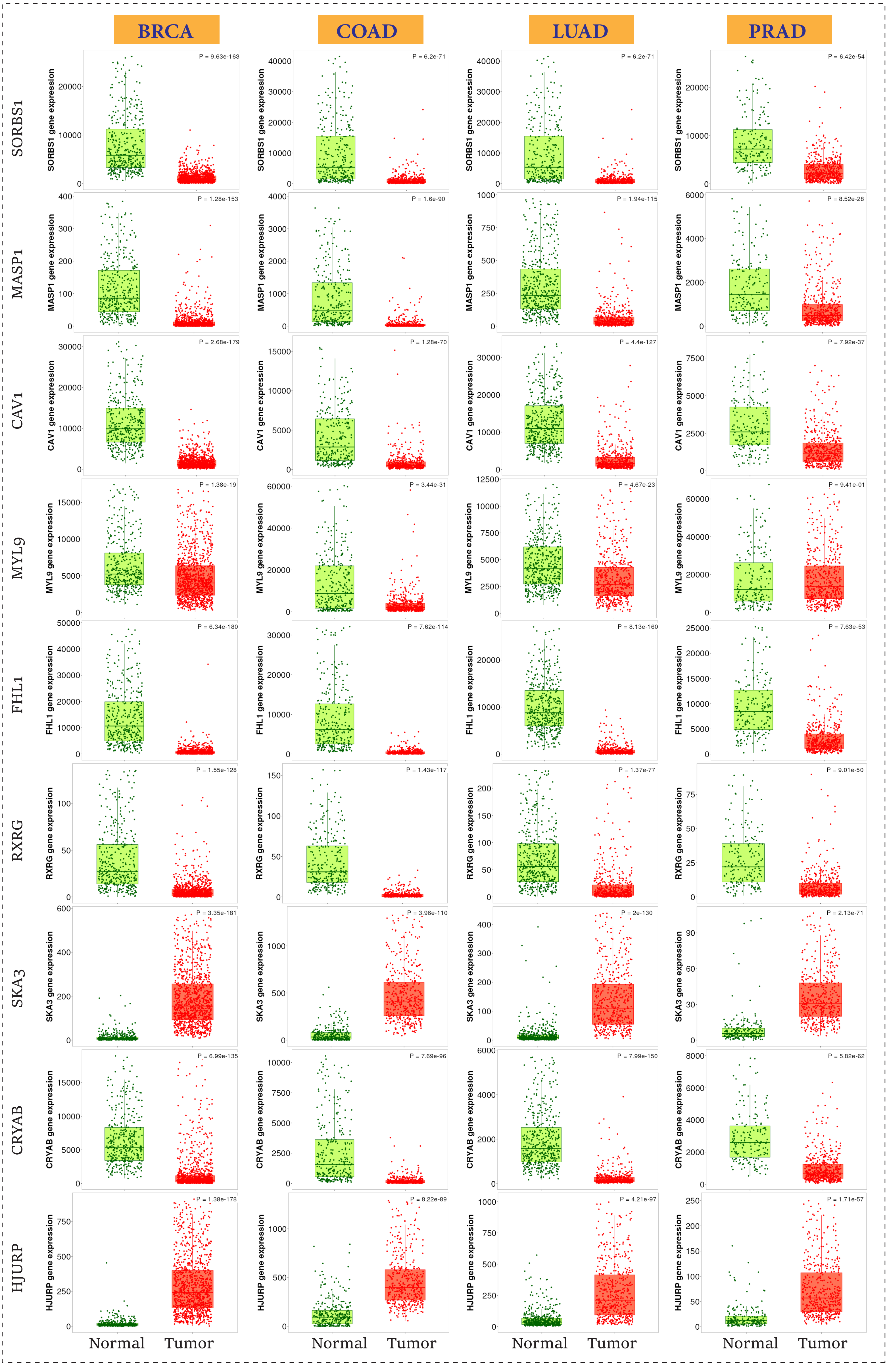

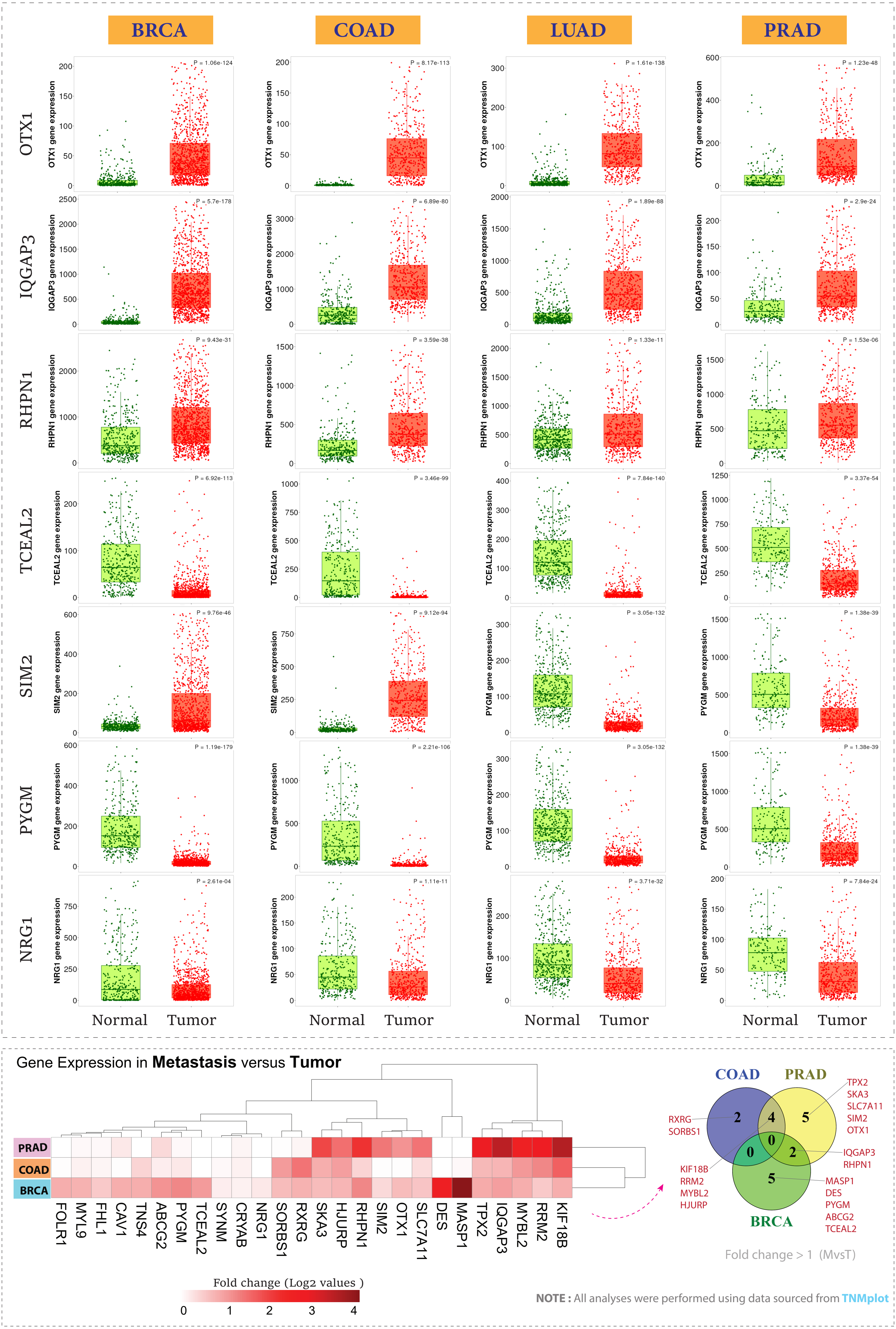

### Suppl. Fig. 5

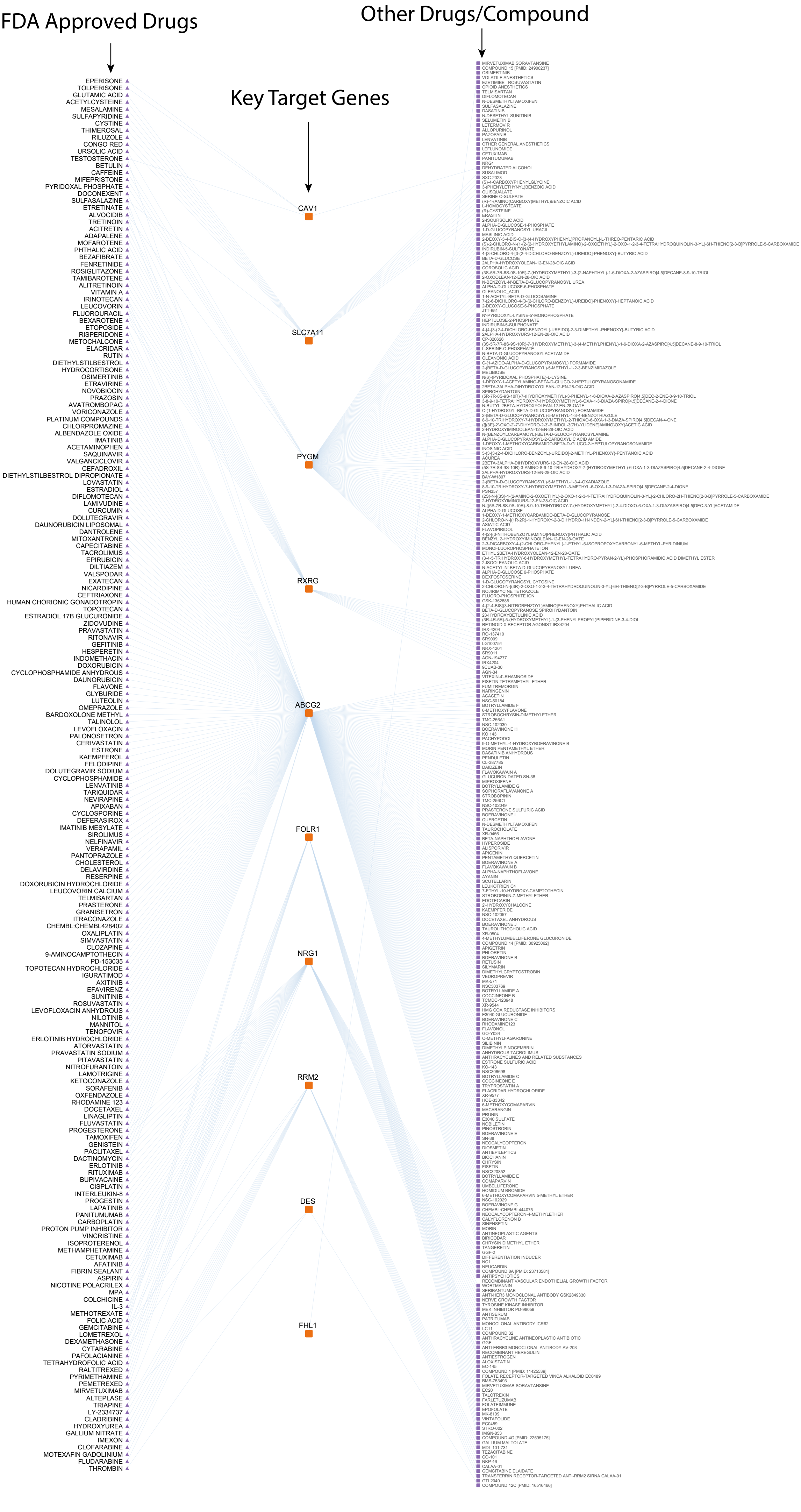

### Suppl. Fig. 6

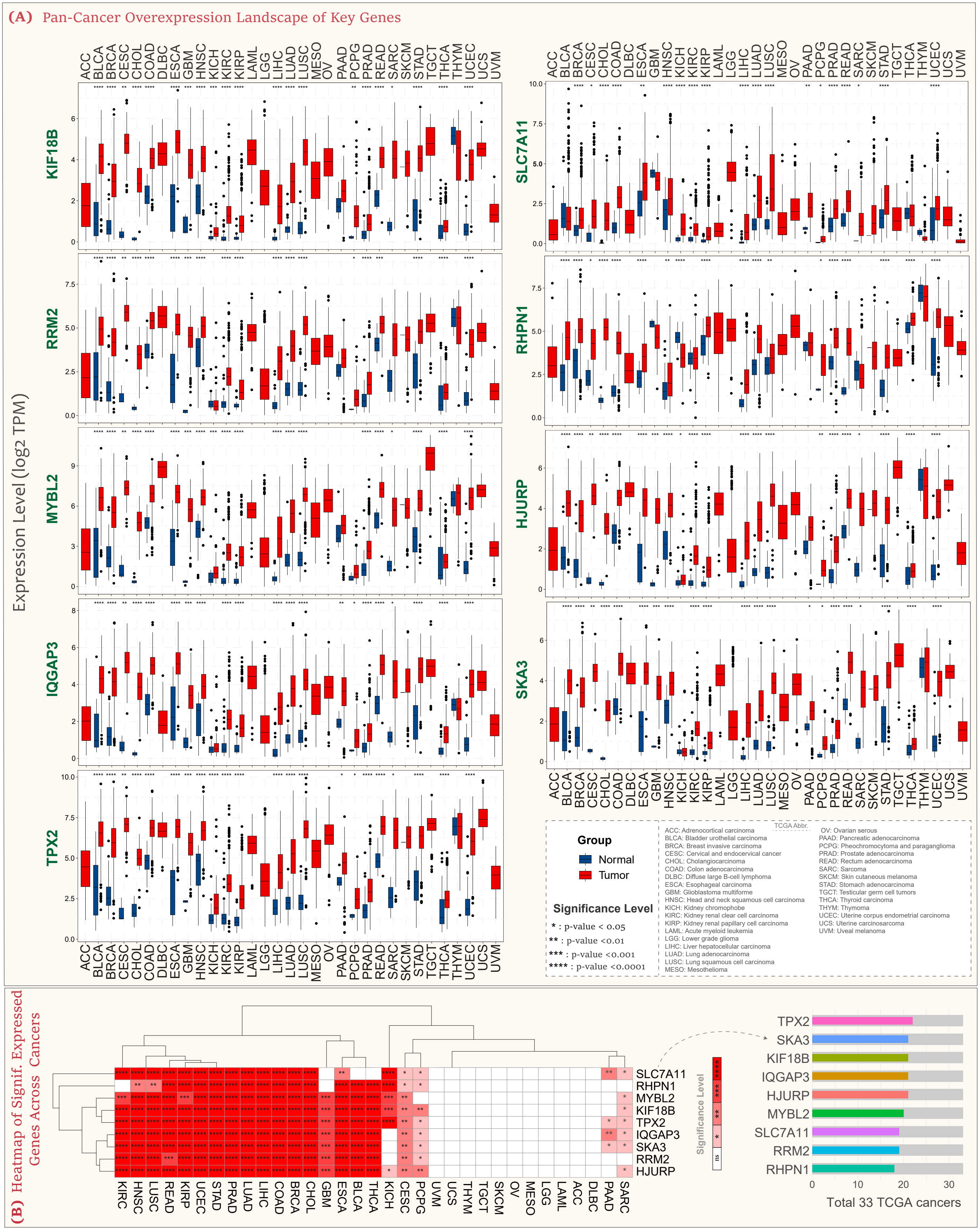
