## Supplementary material for "A Systems Biology Approach to Unveil Shared Therapeutic Targets and Pathological Pathways Across Major Human Cancers": Suppl. Fig. 3

### Hierarchical clustering

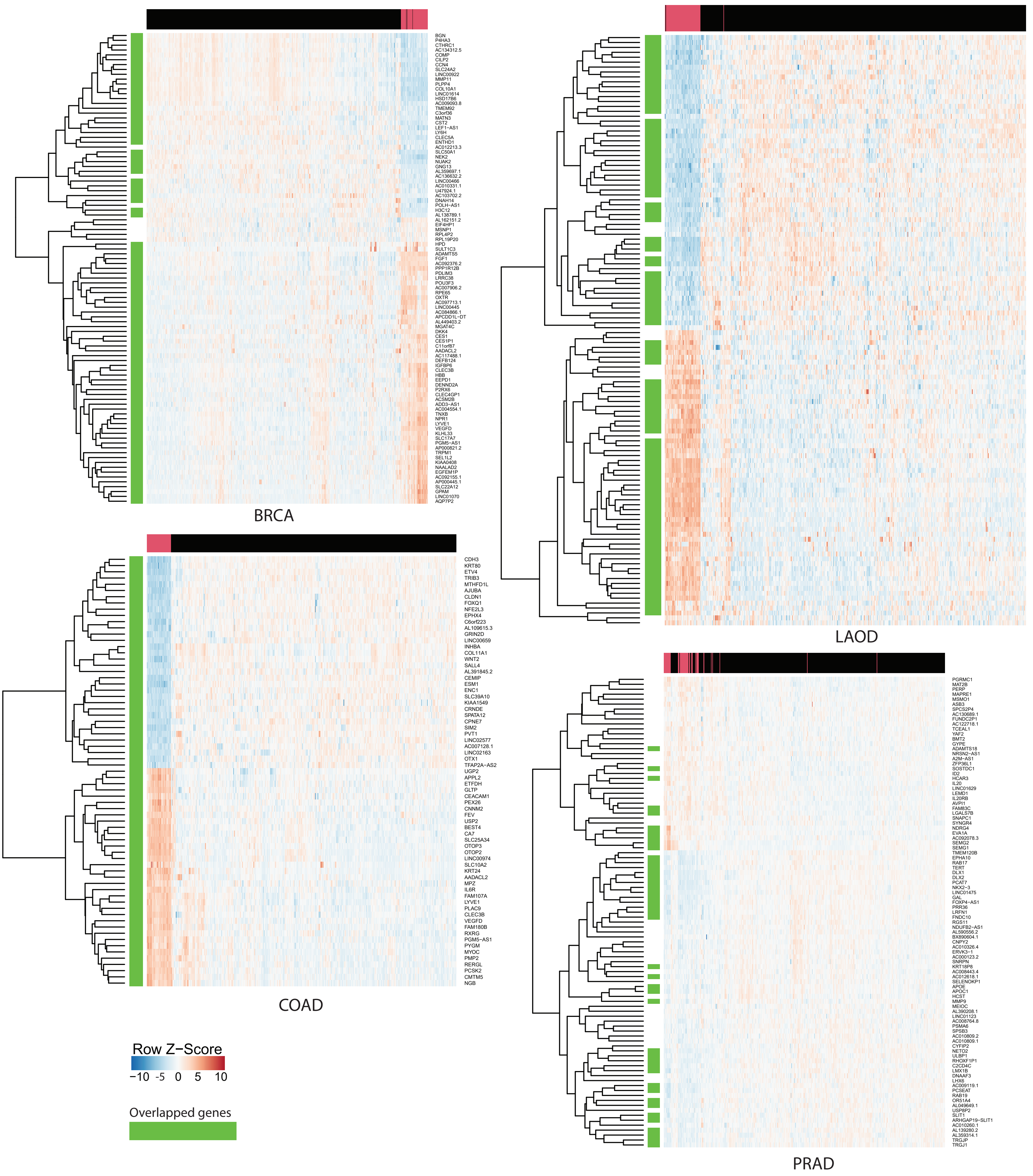


* The genes highlighted in green are the ones that limma had also selected.
* The samples highlighted in red are **Solid Tissue Normal**, the samples highlighted in black are **Primary solid Tumor**.

Please Zoom it.
